# Durable memories and efficient neural coding through mnemonic training using the method of loci

**DOI:** 10.1101/2020.04.29.067561

**Authors:** Isabella C. Wagner, Boris N. Konrad, Philipp Schuster, Sarah Weisig, Dimitris Repantis, Kathrin Ohla, Simone Kühn, Guillén Fernández, Axel Steiger, Claus Lamm, Michael Czisch, Martin Dresler

## Abstract

Mnemonic techniques, such as the method of loci, can powerfully boost memory. Here, we compared memory athletes ranked among the world’s top 50 in memory sports to mnemonics-naïve controls. In a second study, participants completed a six-weeks memory training, working memory training, or no intervention. Behaviorally, memory training enhanced durable, longer-lasting memories. fMRI during encoding and recognition revealed task-based activation decreases in lateral prefrontal, as well as in parahippocampal and retrosplenial cortices in both memory athletes and participants after memory training, partly associated with better performance after four months. This was complemented by hippocampal-neocortical coupling during consolidation, which was stronger the more durable memories participants formed. Our findings are the first to demonstrate that mnemonic training boosts durable memory formation via decreased task-based activation and increased consolidation thereafter. This is in line with conceptual accounts of neural efficiency and highlights a complex interplay of neural processes critical for extraordinary memory.

## Introduction

Mnemonic techniques are powerful tools to enhance memory performance. One of the most common techniques is the so-called “method of loci”, which was developed in ancient Greece and draws upon mental navigation along well-known spatial routes^1^. To-be-remembered material is mentally placed at salient landmarks on an imagined path and can subsequently be recalled by re-tracing the route, “picking up” the previously “dropped” information. The successful application of this method typically requires training and can then lead to exceptional memory performance, as can be seen in individuals participating in events such as the World Memory Championships, and who are able to memorize and accurately reproduce tremendous amounts of arbitrary information (such as word lists, digit series, decks of cards)^2^. In these competitions, however, performance is frequently assessed shortly after study, which makes it impossible to draw conclusions about durable, longer-lasting memories. It is thus unclear whether the method of loci actually helps to form durable rather than weak memories that would eventually fade with time. Previously, we have shown that mnemonics-naïve participants were also able to dramatically boost their memory performance after training the method of loci for several weeks^3^. Here, we drew upon these findings, investigated memory athletes, and tested whether memory training affected durable, longer-lasting memory formation in mnemonics-naïve participants after training.

When applying the method of loci during memory encoding^4–6^ and retrieval^7^, better memory performance appears dovetailed by increased activation within the hippocampus, as well as parahippocampal and retrosplenial cortices. These regions are typically involved in spatial processing^8,9^, including scene construction^10–12^, (mental) navigation^13–15^, and episodic memory^16–19^. Additionally, previous work revealed increased neural processing within the lateral prefrontal cortex when using the technique during encoding^6^, in line with the suggested role of this region in (durable) memory formation^18,20–22^ and in the cognitive control of memory processes via top-down projections^23^. The majority of studies thus far investigated participants who were instructed in the method of loci shortly before the memory tasks were completed (but see^4^). Here, we made a leap forward and performed two separate studies that allowed a detailed characterization of mnemonic expertise, as well as tracking the build-up of experience over time. First, we assessed memory athletes with extraordinary training in using the method of loci, as shown by their ranking among the world’s top 50 in memory sports, and compared them to mnemonics-naïve controls. Second, we recruited mnemonics-naïve participants who underwent either an extensive method of loci training-regime that spanned several weeks, a working memory training, or no intervention. In both studies, we zoomed in onto the neural correlates during encoding and retrieval, and aimed at elucidating their contributions to durable memory formation.

Apart from task-based activation changes, we recently demonstrated the critical role of training-related re-organization of visuo-spatial brain networks during rest, before engaging in any memory-related activity^3^. We found that changes in functional connectivity were associated with increased memory performance in initially naïve participants after method of loci training, becoming similar to those identified in memory athletes. While these training-related alterations were observed during baseline, possibly setting the grounds for optimal memory processing thereafter, it is currently unclear whether memory training also affects connectivity following learning. Such post-task connectivity is thought to reflect consolidation during which memory content becomes stabilized within a wider neocortical network^24,25^. This entails hippocampal-neocortical interactions during awake rest^26,27^ or sleep^28^, potentially indexing the reactivation, or “replay”, of neuronal ensembles that were engaged during the preceding experience^29,30^. In the current work, we investigated hippocampal-neocortical connectivity after learning and its association with durable memory formation after training the method of loci.

Across two separate studies, we tested (1) memory athletes (compared to matched but mnemonics-naïve controls; “athlete study”), and (2) mnemonics-naïve participants who completed an intense method of loci training across six weeks (compared to participants who underwent working memory training or no intervention; “training study”). For all participants, functional magnetic resonance imaging (fMRI) was performed during word list encoding and order recognition, as well as during resting-state periods before and after the tasks. To assess the effects of training, participants of the training study were re-invited to complete another fMRI session after six weeks. Memory performance was assessed during free recall tests immediately and one day after each session (**Fig. 1a-c**). We hypothesized increased memory durability in initially mnemonics-naïve participants after memory training, compared to both control groups. This was expected to be paralleled by training-related neural changes in visuo-spatial brain regions during the tasks, and consolidation-related hippocampal-neocortical coupling during rest. Additionally, we predicted that activation and connectivity profiles in mnemonics-naïve participants after training (compared to the respective control groups) would be similar to when comparing memory athletes with matched controls^3^.

**Fig. 1.:**
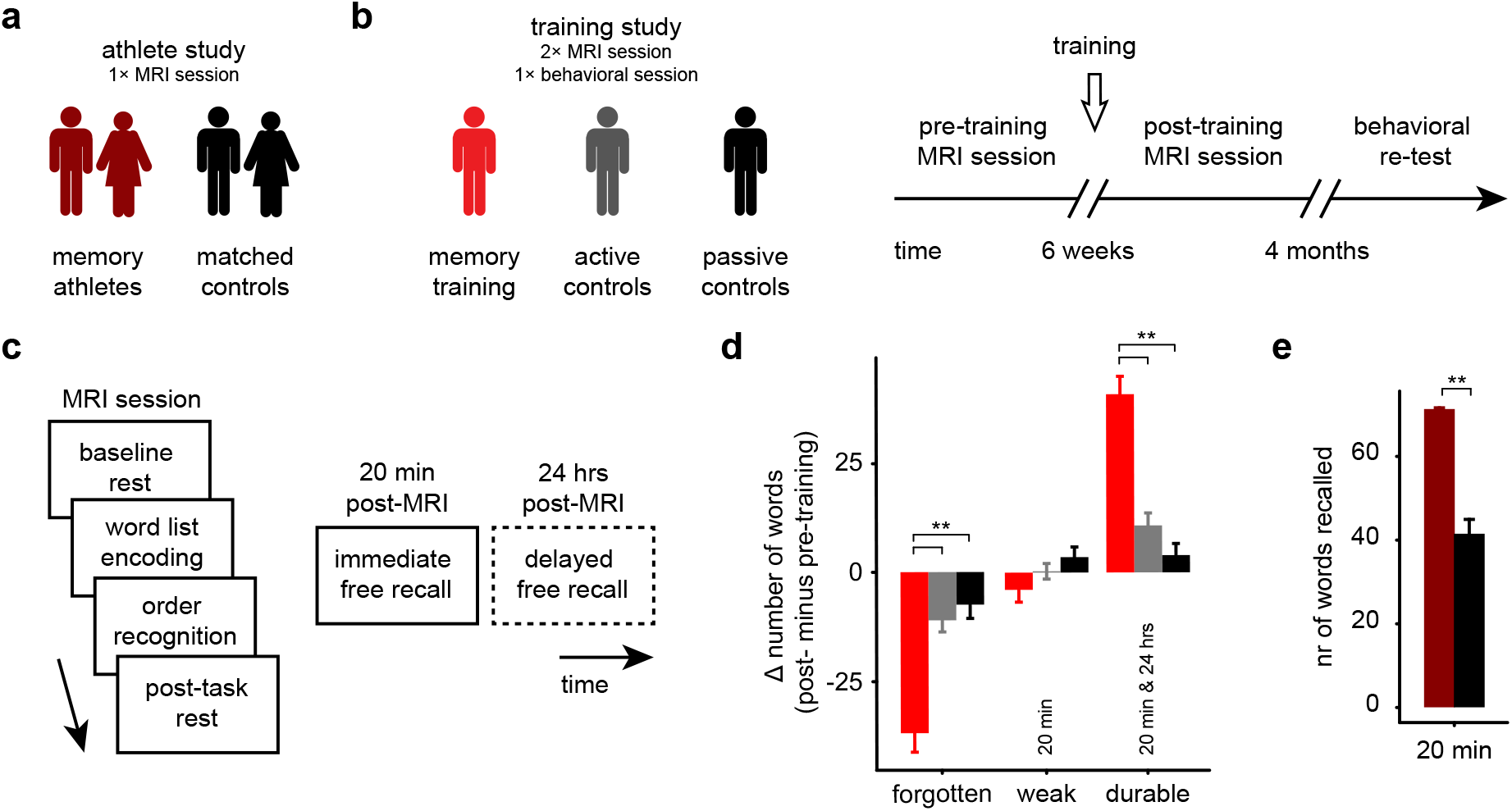
Study design, procedure, and results from the free recall tests. **(a)** In the so-called “athlete study”, we tested memory athletes (*N* = 17) and compared them to matched controls (*N* = 16) during a single MRI session. **(b)** Participants of the “training study” were pseudo-randomized into three groups after an initial MRI session (pre-training): the memory training group (*N* = 17, who underwent method of loci training for six weeks), active controls (*N* = 16, who underwent *n*-back working memory training for six weeks), and passive controls (*N* = 17, who did not participate in any intervention during the six-weeks interval). Participants then returned to the laboratory for a second MRI session (post-training) and took part in a behavioral re-test after four months. **(c)** General structure of MRI sessions: Participants first completed a baseline resting-state period (8 min), the word list encoding (10 min) and order recognition tasks (10 min), as well as a post-task resting-state period (8 min). This was followed by a free recall test outside the MR scanner (5+5 min, 20 min post-MRI, immediate free recall), and another free recall test that was completed from home via phone interview after 24 hours (5+5 min, delayed free recall). The delayed free recall test was only completed by participants of the training study (dashed frame). **(d)** Training study: change in the number of forgotten/weak/durable words from pre- to post-training sessions. Weak memories were only remembered at the immediate free recall test (20 min), while durable memories were also recalled after one day (20 min & 24 hrs). Memory durability during pre- and post-training sessions is shown in **Supplementary Results 1**. ** denotes *p* < 0.0001. **(e)** Athlete study: free recall performance (20 min). ** denotes *p* = 0.0005. Error bars **(d+e)** reflect the standard error of the mean, SEM.

## Results

### Study design and participant samples

We tested 17 participants who were experts in using the method of loci and were ranked among the world’s top 50 in memory sports (hereafter referred to as “memory athletes”), and compared them to a control group closely matched for age, sex, handedness, and intelligence (see also **Methods, Participants of athlete and training studies**). Within this so-called “athlete study”, memory performance and brain function were assessed during a single MRI session (**Fig 1a**; memory athletes: *N* = 17; matched controls: *N* = 16).

In a second study (i.e., the so-called “training study”; *N* = 50), mnemonics-naïve participants completed a method of loci training over six weeks (40 × 30 min). The memory training group (*N* = 17) was compared to an active (*N* = 16) and a passive control group (*N* = 17) that underwent working memory training (40 × 30 min) or no intervention across the six weeks interval, respectively (see **Methods, Study procedures and tasks**). Memory performance and brain function were assessed before and after training, and a behavioral re-test was completed after four months (**Fig 1b**).

### Memory training enhances durable memory formation in initially mnemonics-naïve participants

Starting out, we put our focus on data from the training study and analyzed the change in memory durability from before to after training (for a discussion of overall free recall performance, see^3^). During both sessions, participants studied word lists and were asked to retrieve material during a free recall test 20 min after MR scanning (immediate free recall), as well as 24 hours later (delayed free recall; **Fig 1c**). While some words were never recalled (i.e., forgotten material), weak memories were defined as those that could only be remembered at the immediate free recall but were forgotten afterwards. Durable memories were the ones also remembered after the delay.

Results revealed a significant increase in durable memories, along with decreased forgetting, in the memory training group from before to after training, compared to both active and passive control groups (**Fig. 1d**). The change in the amount of weak memories from pre- to post-training did not significantly differ between the three groups (mixed-model analysis of variance (ANOVA); memory training group, *N* = 17; active controls, *N* = 16; passive controls, *N* = 17; group × memory type interaction, *F*(4,141) = 32.83, *p* < 0.0001; main effect of group, *p* > 0.999; main effect of memory type, *F*(2,141) = 13.27, *p* < 0.0001; see **Supplementary Table S1** for a list of all significant pair-wise comparisons). Additional analyses revealed that the change in memory durability was specifically related to memory training and was not due to potential performance differences already present pre-training (**Supplementary Results 1**). Thus, training the method of loci increased durable memories in initially mnemonics-naïve participants.

Different from the training study, the athlete study comprised a single experimental session; performance was tested immediately after the tasks and only once (20 min post-MRI; thus, the analysis of memory durability was not possible). As also shown previously^3^ (but note that current analyses involved a sub-sample of participants), free recall performance within the athlete study was generally high, and was significantly higher in memory athletes compared to matched controls (Wilcoxon signed-rank test; one matched pair excluded from analysis, memory athletes, *N* = 16, median = 72; matched controls, *N* = 16, median = 43; *V* = 136, *p* = 0.0005; **Fig. 1e**).

### Method of loci decreases activation in lateral prefrontal cortex during word list encoding in athletes and initially mnemonics-naïve participants after training

Next, we turned to the fMRI data and investigated changes in brain activation from pre- to post-training while participants studied novel words (word list encoding task; **Fig. 1c**, **Fig 2a**). Memory athletes and the memory training group (post-training) were asked to use the method of loci during encoding. We thus hypothesized engagement of regions typically involved in visuo-spatial processing and successful memory encoding, including the hippocampus and adjacent MTL structures, retrosplenial cortex, and lateral prefrontal regions^4–6,18,22^. Notably, we refrained from a common subsequent memory analysis as trials were unevenly distributed between groups (e.g., most memory athletes might remember all words and forget none, whereas this might be different for participants of the control groups). We thus opted for an alternative approach and tested activation changes during encoding compared to an implicit baseline, with individual memory durability scores added as a covariate (see also **Methods, MRI data processing: task data**).

**Fig. 2.:**
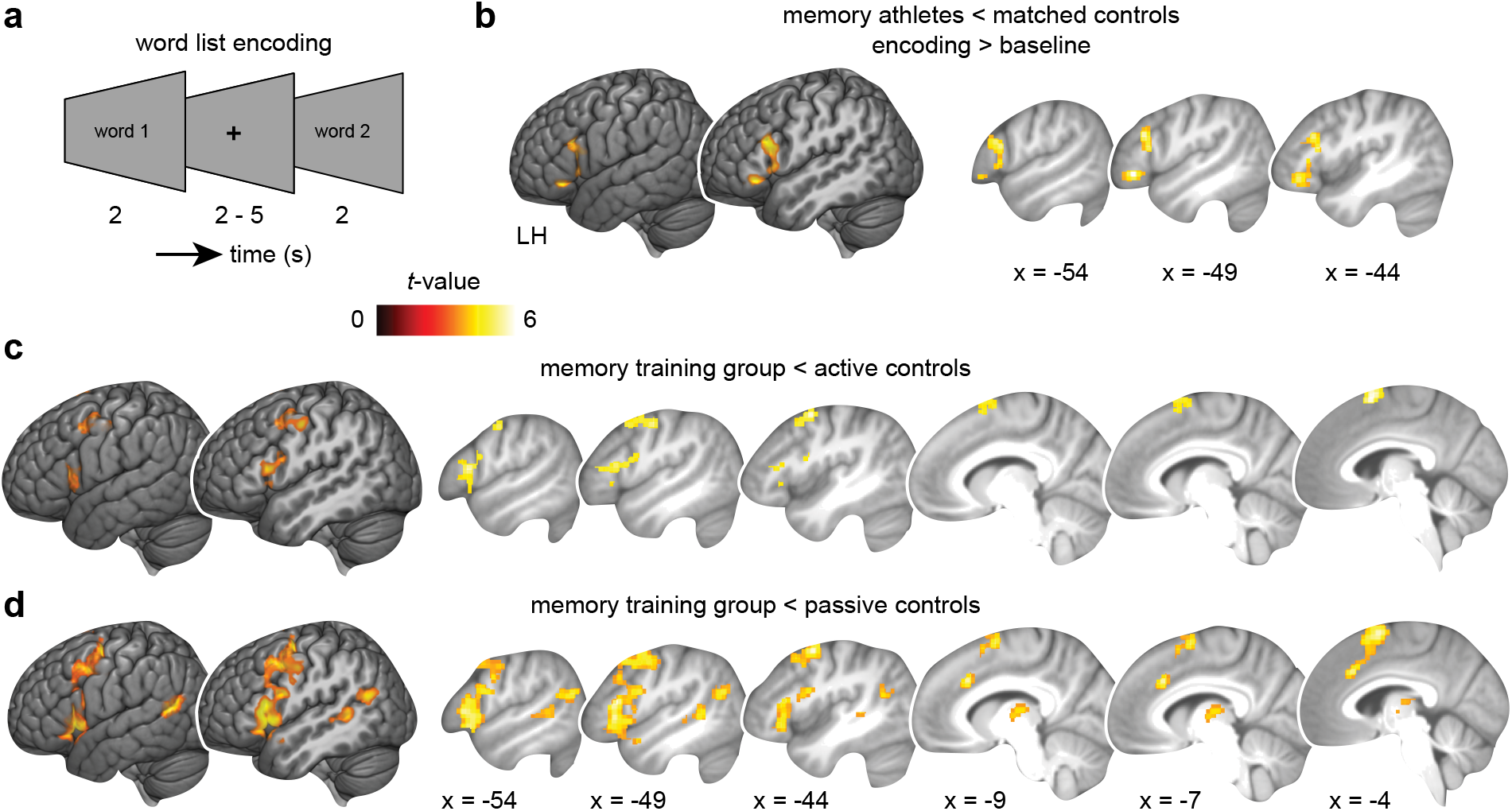
fMRI results from word list encoding. **(a)** Word list encoding task: Participants studied novel words during each MRI session. **(b)** Athlete study: brain activation during encoding (encoding > baseline) is decreased in memory athletes compared to matched controls (see also **Supplementary Table S2**). **(c+d)** Training study: brain activation is significantly decreased in the memory training group after training when comparing to **(c)** active or **(d)** passive controls (group × session interactions; see **Supplementary Tables S3** and **S4**, also for main effects; and **Supplementary Table S5** for a comparison between active and passive controls). Results are shown at *p* < 0.05 family-wise error (FWE)-corrected at cluster level (cluster-defining threshold *p* < 0.001). LH, left hemisphere.

First, we focused on data from the athlete study and compared memory athletes to matched controls. Surprisingly, we found robust activation *decreases* within the left lateral prefrontal cortex during word list encoding (independent-samples *t*-test, contrast encoding > baseline, covariate number of words freely recalled, memory athletes, *N* = 17; matched controls, *N* = 16, statistical threshold for this and all following analyses: *p* < 0.05, family-wise error (FWE) corrected at cluster level using a cluster-defining threshold of *p* < 0.001; **Fig. 2b**; **Supplementary Table S2**). Contrary to what we had expected, there were no significant activation changes within the MTL or retrosplenial cortex.

Second, we leveraged data from the training study and found, strikingly similar to above, decreased activation within the left lateral prefrontal cortex in the memory training group after training, compared to both the active and the passive control groups (interaction effects, two separate full factorial designs, contrast encoding > baseline, covariate memory durability score, memory training group, *N* = 17; active controls, *N* = 16; passive controls, *N* = 17; **Fig. 2cd**; **Supplementary Tables S3** and **S4**, also for main effects of group and session). When comparing the memory training with the passive control group, activation decreases further included the thalamus and the left angular gyrus (**Fig. 2cd**, **Supplementary Table S4**; and see **Table S5** for a comparison between active and passive control groups). Thus, results from both studies consistently revealed decreased activation within left lateral prefrontal regions when applying the method of loci during word list encoding.

To elucidate if results were actually caused by a decrease in activation in the memory training group over time rather than by group differences already present pre-training, we performed additional region-of-interest (ROI) analyses and extracted average activation values from significant interaction clusters, altogether confirming the results above (**Supplementary Results 2**). Finally, there were no significant differences in activation levels between the memory athletes and the memory training group (post-training) during word list encoding (independent-samples *t*-test, contrast encoding > baseline, covariate number of words recalled during the immediate free recall test, memory athletes, *N* = 17; memory group (post-training), *N* = 17), indicating similar activation profiles of athletes and initially mnemonics-naïve participants after training the method of loci.

### Expertise in the method of loci can positively affect recognition performance while slowing down response times

Following word list encoding, participants completed the order recognition task where word triplets were presented in either the same or a different order as studied previously (**Fig. 1c**, **Fig. 3a**). We expected memory athletes and participants in the memory training group (post-training) to excel at this task as they would utilize the method of loci to more efficiently retrieve the order of the presented words.

**Fig. 3.:**
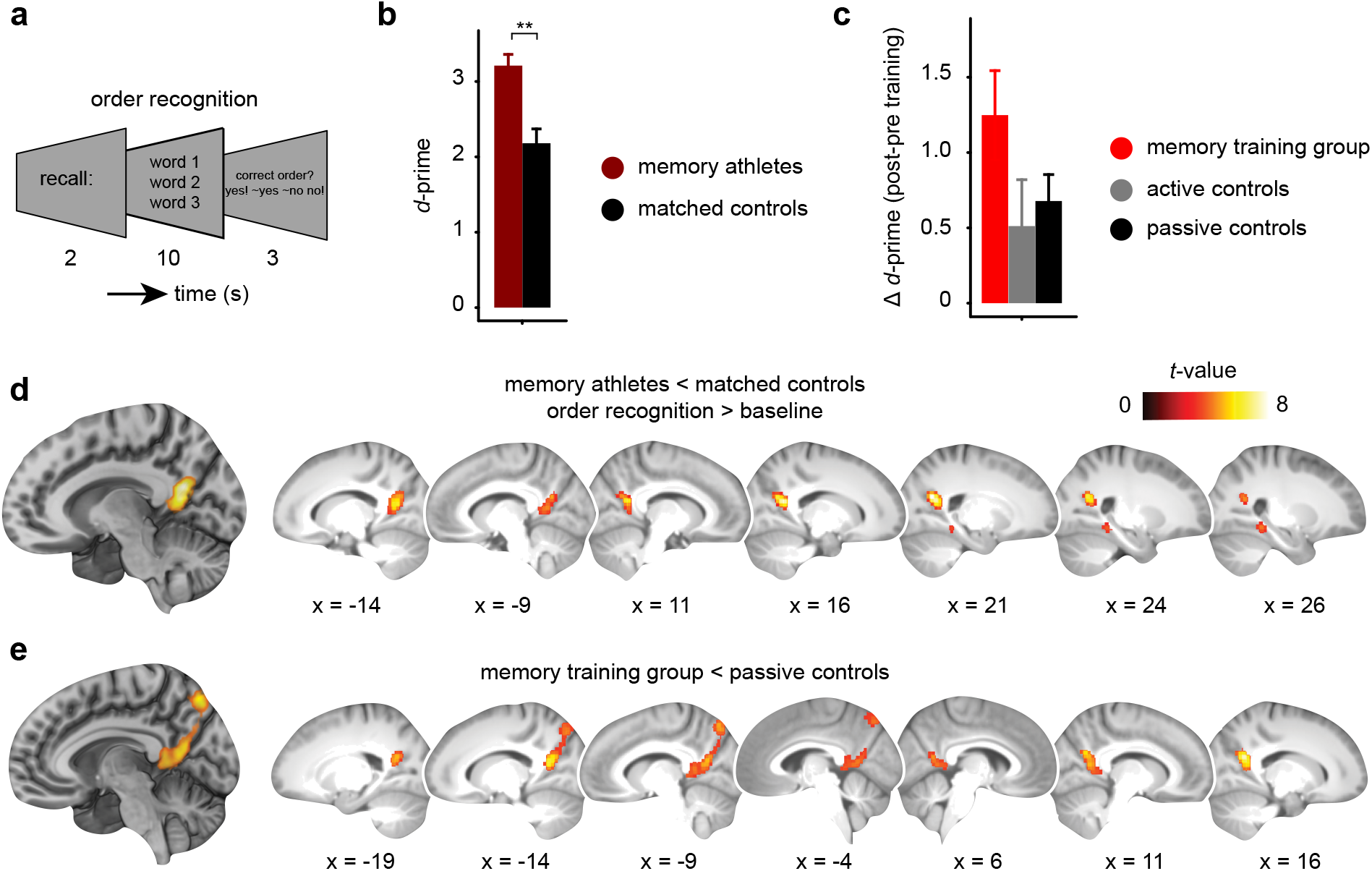
Behavioral and fMRI results from order recognition. **(a)** Order recognition task: after word list encoding, word triplets were presented in the same or a different order as studied previously and participants were asked to judge the order. **(b)** Athlete study: recognition performance (*d*-prime) for memory athletes and matched controls. ** denotes *p* < 0.001. Results for response times (RTs) are presented in **Supplementary Results 3**. **(c)** Training study: change in *d*-prime (from pre- to post-training sessions) across the groups (main effect of group, *p* = 0.133). Recognition performance during pre- and post-training sessions is shown in **Supplementary Results 4**. Results for response times (RTs) are presented in **Supplementary Results 3**. Error bars **(b+c)** reflect the standard error of the mean, SEM. **(d)** Athlete study: brain activation during order recognition (order recognition > baseline) is decreased in memory athletes compared to matched controls (see also **Supplementary Table S6**). **(e)** Training study: brain activation is significantly decreased in the memory training group after training when comparing to passive controls (group × session interaction; see **Supplementary Table S7**, also for main effects; and see **Supplementary Table S8** for a comparison between memory training group and active controls). Results are shown at *p* < 0.05 FWE-corrected at cluster level (cluster-defining threshold *p* < 0.001).

Memory athletes indeed showed significantly higher recognition performance (indexed through *d*-prime) compared to matched controls (**Fig. 3b**; independent-samples *t*-test; memory athletes, *N* = 17; matched controls, *N* = 16; *t*(30.2) = 4.27, *p* < 0.001; and see **Supplementary Results 3** for response times). However, despite numerically increased *d*-prime scores in the memory training group after training, there was no significant difference in recognition performance between the three groups (change in *d*-prime; **Fig. 3c**; one-way ANOVA; memory training group, *N* = 17; active controls, *N* = 16; passive controls, *N* = 17; main effect of group, *p* = 0.133, pair-wise comparisons: memory training > active controls: *p* = 0.135, effect size *d* = 0.592; memory training > passive controls: *p* = 0.294, *d* = 0.569; active > passive controls: *p* = 0.898, *d* = −0.161; see **Supplementary Results 4** for a general increase in *d*-prime from pre- and post-training). Additionally, participants of the memory training group showed slower response times after training (**Supplementary Results 3**). Thus, training the method of loci increased recognition performance in memory athletes. While the effect of memory training on recognition performance in initially mnemonics-naïve participants appeared to be positive as well (but note that results were not significant), results were accompanied by generally slower responses.

### Method of loci decreases activation in posterior parahippocampal and retrosplenial cortices during order recognition in athletes and initially mnemonics-naïve participants after training

We next turned to the neuroimaging data acquired during order recognition. Since memory athletes and participants of the memory training group (post-training) were asked to use the method of loci during order recognition, we expected increased engagement of brain regions typically associated with visuo-spatial processing and successful memory retrieval, such as the hippocampus, parahippocampal and retrosplenial cortices^4,7–9,13,16,19^. Again, as correct and incorrect trials were unevenly distributed between groups, we compared recognition trials against the implicit baseline, with individual *d*-prime scores added as a covariate (see also **Methods, MRI data processing: task data**).

Similar to the profile of activation decreases during the preceding word list encoding task (see above), results indicated reduced activation within the right posterior parahippocampal and bilateral retrosplenial cortices in memory athletes compared to matched controls (independent-samples *t*-test, contrast order recognition > baseline, covariate *d*-prime, memory athletes, *N* = 17; matched controls, *N* = 16, **Fig. 3d**, **Supplementary Table S6**). This was dovetailed by decreased activation within the posterior parahippocampal and bilateral retrosplenial cortices, and in the precuneus in the memory training group after training, when comparing to passive controls (interaction effect, full factorial design, contrast order recognition > baseline, covariate *d*-prime, memory training group, *N* = 17; passive controls, *N* = 17; **Fig. 3e**; and see **Supplementary Table S7** for main effects of group and session). There was no significant group × session interaction when comparing the memory training group with active controls, but activation in precuneus and superior parietal gyri generally decreased over time (main effect of session, **Supplementary Table S8**, full factorial design, contrast order recognition > baseline, covariate *d*-prime, memory training group, *N* = 17; active controls, *N* = 16). Furthermore, active and passive control groups did not differ significantly (full factorial design, contrast order recognition > baseline, covariate *d*-prime, active controls, *N* = 16; passive controls, *N* = 17). To summarize, findings from both studies consistently revealed decreased activation within the posterior parahippocampal and retrosplenial cortices when applying the method of loci during order recognition.

Additional ROI-analyses confirmed that these results were actually related to memory training and not an effect of potential group differences already present pre-training (**Supplementary Results 5**). Finally, as also for data during word list encoding (see above), there were no significant differences in activation levels between the memory athletes and the memory training group (post-training) during order recognition (independent-samples *t*-test, contrast order recognition > baseline, covariate *d*-prime, memory athletes, *N* = 17; memory group (post-training), *N* = 17), indicating similar activation profiles in athletes and initially mnemonics-naïve participants after training the method of loci.

### Training-related activation decreases during order recognition are associated with better memory performance during the four-months re-test

Next, we asked whether the whole-brain activation changes from pre- to post-training sessions during the memory tasks (word list encoding, order recognition) were associated with increased memory performance beyond the 24-hour delay. During the re-test after four months, participants of the training study were once more invited to the behavioral laboratory where they completed the word list encoding task followed by a free recall test (the same word list as during the pre-training session was used; **Methods, Study procedures and tasks, Re-test after 4 months**).

As also reported previously^3^ (but note that current analyses include a sub-sample of participants), the memory training group showed significantly increased memory performance after four months (free recall; 4-month re-test minus pre-training test 20 min post-MRI) as compared to both active and passive control groups (change in the number of words freely recalled, mean ± SEM: memory training group: 22.67 ± 4.87; active controls: −0.71 ± 2.65; passive controls: −1.5 ± 2.79; five subjects were not available for the re-test, analysis thus included 45 participants; memory training group, *N* = 16; active controls, *N* = 14; passive controls, *N* = 15; one-way ANOVA; main effect of group, *F*(2,42) = 14.67, *p* < 0.0001; pair-wise comparisons: memory training > active controls, *t*(42) = 4.53, *p* = 0.0001; memory training > passive controls, *t*(42) = 4.84, *p* = 0.0001; active controls > passive controls, *p* = 0.987). Hence, the memory training group was able to utilize the method of loci successfully (as indicated through increased memory performance compared to both control groups), even after several months.

We then went on to test the cross-participant relationship between whole-brain activation decreases from pre- to post-training and the change in memory performance (4-month re-test minus pre-training_20min_). To this end, we created individual difference maps (pre-minus post-training) based on the first-level contrasts (encoding > baseline, order recognition > baseline), reflecting decreased activation over time. These difference maps were then submitted to two separate linear regression analyses with the change in memory performance added as a covariate of interest.

During order recognition, activation decreases across sessions were positively associated with memory performance after four months. This included decreased activation from pre- to post-training within a widespread set of regions comprising the hippocampus, the posterior parahippocampal region, the left fusiform gyrus, retrosplenial cortex, precuneus, left angular gyrus, thalamus, bilateral striatum, medial prefrontal and orbitofrontal cortex, and the precentral gyrus (**Fig. 4**, **Supplementary Table S9**). Thus, stronger activation decreases in these regions during order recognition were coupled to increased memory performance at the four-months re-test across all participants of the training study. In contrast, activation decreases during word list encoding appeared unrelated to memory performance after four months.

**Fig. 4.:**
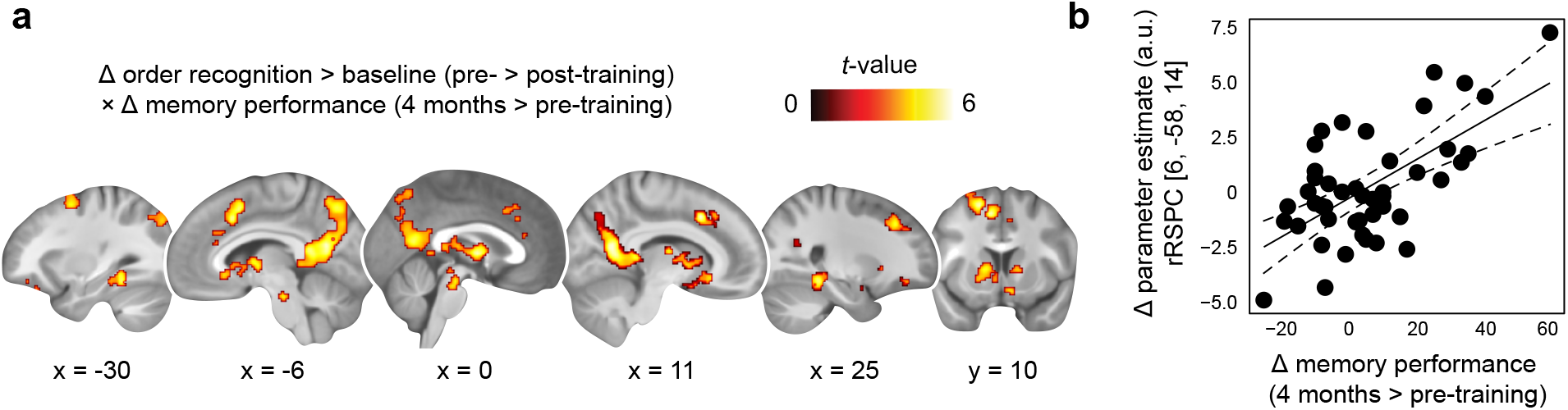
Activation decreases during order recognition and association with memory performance during the 4-months re-test. **(a)** Training study: decreases in brain activation (order recognition > baseline) from before to after training (pre- > post-training) that positively scaled with the change in memory performance from the pre-training session (20 min post-MRI) to the re-test after four months (covariate of interest). Results are shown at *p* < 0.05 FWE-corrected at cluster level (cluster-defining threshold *p* < 0.001; see also **Supplementary Table S9**). **(b)** For visualization, the scatter plot shows the relationship between the change in parameter estimates (arbitrary units, a.u.) from the pre- to post-training sessions, extracted from the global maximum (right retrosplenial cortex, rRSPC, 8 mm sphere around MNI peak coordinate, x = 6, y = −58, z = 14), and the change in memory performance (4-months re-test minus pre-training_20min_).

### Increased hippocampal-neocortical coupling during post-task rest is related to memory consolidation in athletes and initially mnemonics-naïve participants after training

So far, we documented increased memory performance (memory durability, recognition performance), along with decreased brain activation during memory-related processing (word list encoding, order recognition) in memory athletes and participants of the memory training group after training. Additionally, we hypothesized that durable memory formation should be associated with increased consolidation processes during rest after learning, involving hippocampal-neocortical circuits^25–27,31^. To test this, we took the anatomical boundaries of the bilateral hippocampus as a seed (**Fig. 5a**), calculated its whole-brain connectivity during each resting-state period and session, and tested if connectivity varied as a function of memory performance (i.e., free recall performance in the athlete study, memory durability in the training study; see also **Methods, MRI data processing: resting-state periods**).

**Fig. 5.:**
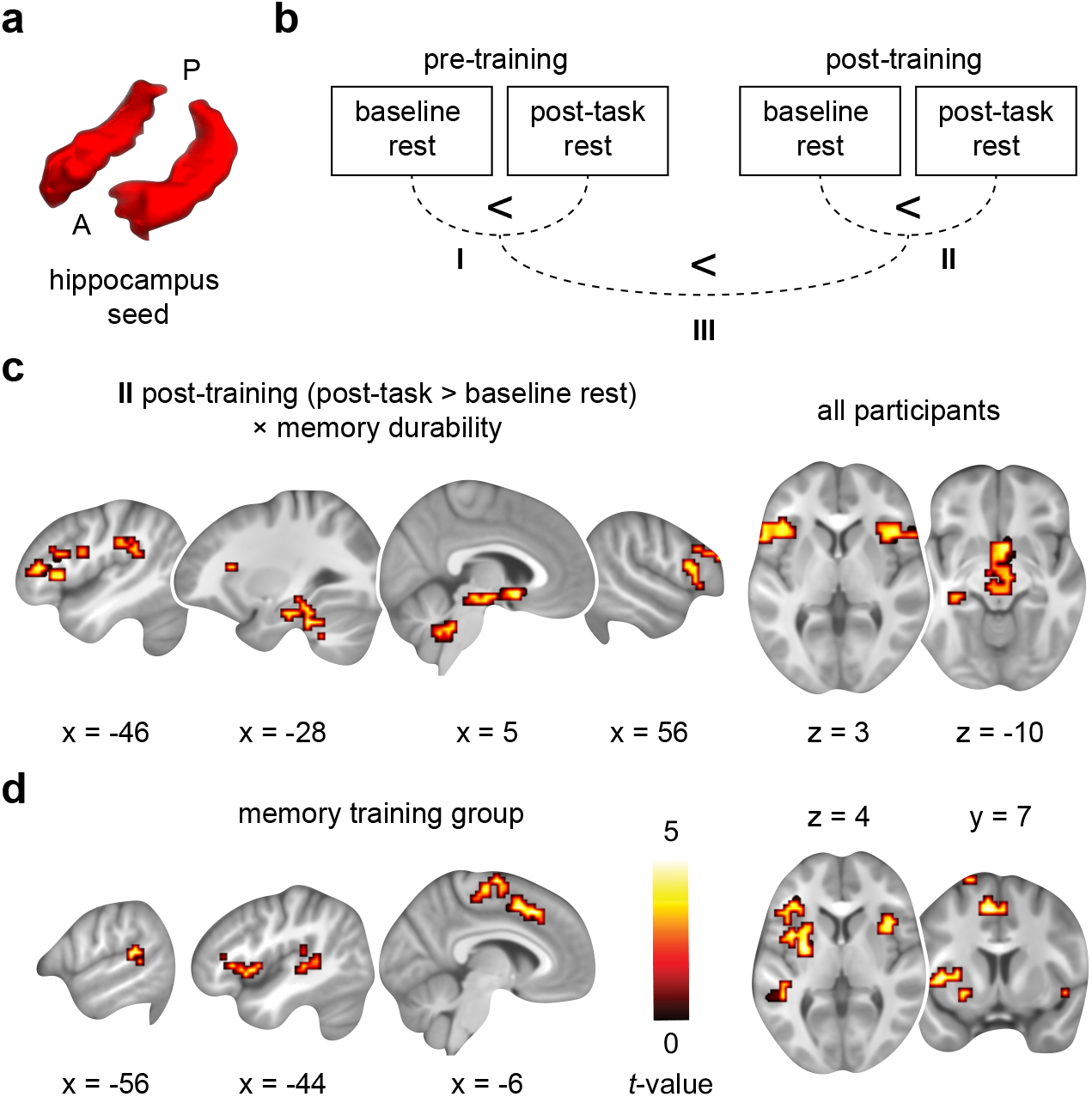
Increased hippocampal-neocortical coupling during post-training rest and relationship with durable memory formation. **(a)** Bilateral anatomical hippocampus seed used for whole-brain connectivity analysis. A, anterior, P, posterior. **(b)** Training study: schematic of the analysis steps performed. We tested hippocampal connectivity increases (post-task > baseline rest) during the pre- (**I**) and post-training sessions (**II**), and investigated the increase in consolidation-related coupling from pre- to post-training sessions (**III**, [post-task > baseline rest]_post_ > [post-task > baseline rest]_pre_). Analysis of data from the athlete study involved a single MRI session (post-task > baseline), which is not depicted here. **(c)** Training study: hippocampal-neocortical connectivity increases from baseline to post-task rest during the post-training session positively scaled with the proportion of durable memories formed (i.e., memory durability) across all participants (see also **Supplementary Table S10**). **(d)** Follow-up analyses revealed that these effects were specifically driven by connectivity changes in the memory training group, but were not present in passive or active controls (see **Supplementary Results 7**, and **Supplementary Table S11**). All results are shown at *p* < 0.05 FWE-corrected at cluster level (cluster-defining threshold *p* < 0.001).

First, we focused on data from the athlete study and investigated consolidation-related hippocampal connectivity increases from before to after the tasks. Across participants (including both memory athletes and matched controls, *N* = 33), we found coupling between the hippocampus and a bilateral cerebellar region that positively scaled with subsequent free recall performance (linear regression, contrast difference map (post-task > baseline rest), number of words freely recalled 20 min post-MRI added as a covariate of interest; *p* < 0.05, FWE-corrected at cluster level using a cluster-defining threshold of *p* < 0.001, cluster size = 62 voxels; MNI peak coordinate of global maximum, x = 32, y = −70, z = −52; *Z*-value = 4.6, 416 voxels). Thus, hippocampal connectivity with the cerebellum was stronger during rest the more words participants recalled. More specifically, these results appeared to be driven by stronger hippocampal-cerebellar coupling in memory athletes compared to matched controls (**Supplementary Results 6**).

Second, we turned towards data from the training study (including the memory training group, active and passive controls, *N* = 49). To draw precise conclusions about connectivity changes related to extensive memory training, we investigated hippocampal-neocortical coupling before and after training, as well as changes from before to after training and its association with durable memory formation. Correspondingly, this involved three analysis steps: connectivity (I) during the pre-training session (post-task > baseline rest), (II) during the post-training session (post-task > baseline rest), and (III) changes from pre- to post-training sessions ([post-task > baseline rest]_post_ > [post-task > baseline rest]_pre_; see also **Fig. 5b**).

Results from the post-training session (II; **Fig. 5bc**) revealed stronger connectivity from baseline to post-task rest between the hippocampus and the bilateral lateral prefrontal cortex, left angular gyrus, the left hippocampus and parahippocampal cortex, bilateral insula and right caudate nucleus, as well as the brainstem and cerebellum that positively scaled with memory durability across participants (linear regression, contrast difference map (post-task > baseline rest), memory durability (post-training) added as a covariate of interest; see **Fig. 5c**). There was no negative association between hippocampal-neocortical connectivity and memory durability (but see **Supplementary Table S10** for general connectivity increases from baseline to post-task rest). Therefore, hippocampal coupling with a widespread set of neocortical regions was stronger after training the more durable memories participants formed. Follow-up analyses confirmed that this effect was driven by connectivity changes in the memory training group and was not present in the active or passive controls (**Fig. 5d**; and see **Supplementary Results 7**).

We did not find any significant, hippocampal-neocortical connectivity increases related to memory durability during the pre-training session (I; **Fig. 5b**), or from pre- to post-training sessions (III; **Fig. 5b**; but see **Supplementary Table S12** for general connectivity increases across sessions), and none of the results were associated to the change in memory performance from pre-training to after four months.

To summarize, stronger hippocampal-cerebellar connectivity during rest after memory processing was associated with increased memory performance across participants of the athlete study. In the training study, hippocampal connectivity with the lateral prefrontal cortex, MTL, and striatum was increased the more durable memories participants formed (post-training). Additional analyses confirmed that these effects were specifically driven by connectivity changes in memory athletes and the memory training group after training, but were not present in any of the control groups.

## Discussion

In this study, we investigated memory training using the method of loci, and its impact on memory durability and neural coding. To obtain a detailed characterization of long-term training effects and existing expertise with mnemonic techniques, we performed two separate studies that involved memory athletes as well as mnemonics-naïve participants who underwent an extensive memory training over six weeks. Most importantly, and in line with theoretical accounts of neural efficiency, memory training enhanced durable memories and decreased task-based activation while increasing hippocampal-neocortical consolidation.

Central to our question was the potential effect of mnemonic training on durable memory formation. We found that initially naïve participants improved memory durability after training, compared to both active and passive control groups (**Fig. 1d**). These results were mirrored by the exceptional, close-to-ceiling performance in memory athletes compared to matched controls (**Fig. 1e**). Effectively using the method of loci requires mental navigation along well-known spatial routes and the anchoring of to-be-remembered information to salient locations on the path^1^. The method thus combines several key aspects that are thought to affect memory. First, the method of loci relies on visuo-spatial processing that engages the hippocampus, parahippocampal and retrosplenial cortices^4–7^. These brain regions are typically associated with spatial processing and (mental) navigation^8–15,32^, as well as (episodic) memory^16–19,32^. A link between space and memory therefore appears natural, and spatial representations have been discussed to organize conceptual knowledge and to allow flexible behavior^13,33^. Second, the reliance on well-known spatial routes bears resemblance to the utilization of schema-like knowledge structures that are established during prior experiences^34^. Schemas are assumed to provide a scaffolding to promote memory encoding and consolidation^35^. Instinctively, the stable formation of spatial routes for mental navigation takes time and should thus benefit from extensive method of loci training. While previous studies provided participants with an introduction into the mnemonic technique one day prior^5^ or shortly before study^7^, we recruited participants who underwent a training-regime that spanned several weeks (see also^3^). Hence, our training allowed participants to build-up stable spatial routes that could incorporate novel information more readily, drastically enhancing durable memories and sustainably increasing performance even after four months. Related to this, mnemonic techniques were discussed to speed-up memory stabilization using schema-like structures, promoting the direct transfer from working memory into long-term storage (as proposed by the “long-term working memory” hypothesis^36^). Third, mentally placing arbitrary to-be-remembered information at salient locations along the imagined path likely produces relatively bizarre associations, thereby triggering neural mechanisms related to novelty^35,37^. This in turn cranks up dopaminergic and noradrenergic release from the brainstem and ventral striatum towards the hippocampus^38,39^, which is thought to promote memory persistence by triggering synaptic^40^ and systems consolidation^25^. Overall, we suggest that the method of loci favorably combines the abovementioned aspects (visuo-spatial processing, prior knowledge, novelty) to boost durable memories, leading to exceptional memory performance in athletes and initially mnemonics-naïve participants after training.

Memory formation is thought to rely on successful encoding. We found consistent activation decreases in lateral prefrontal regions when memory athletes and participants of the memory training group (post-training) studied verbal material (**Fig. 2b-d**). The lateral prefrontal cortex is involved in memory encoding while applying the method of loci^6^, supports durable memory formation^18,20–22^, the selection and organization of memories via top-down control^23^, and allows flexible behavior^41^. Decreased activation in lateral prefrontal regions might hence indicate a diminished requirement for cognitive control due to extensive method of loci training. Similarly, we found decreased activation within the posterior parahippocampal and retrosplenial cortices during order recognition in memory athletes and participants of the memory training group after training (**Fig. 3de**). Although these results are similar to previous reports with regards to their spatial layout^4–7^, we revealed diametrically opposite effects. In other words, we report robust activation *decreases* despite that successful memory encoding^18,20–22^ and retrieval^16,19,42^ typically engage *increased* activation in a set of prefrontal, medial temporal, and visuo-spatial brain regions. Importantly, the training-related decreases were directly associated with better memory performance at the four-months re-test across participants (**Fig. 4**), mainly attributable to connectivity changes in the memory training group after training.

Our results are in line with the so-called “neural efficiency hypothesis”^43^, which proposes that highly-skilled or intelligent individuals display lower (thus, more efficient) brain activation during cognitive tasks for reaching the same behavioral performance^44–48^. For instance, participants with higher verbal or visuo-spatial skills were found to show lower levels of brain activation when using the respective strategies during cognitive tasks^45^. Such efficient neural coding might require extensive training^43^. Our six-weeks training-regime might thus resemble the build-up of expertise and could explain the differential findings compared to previous studies. The concept of neural efficiency has, however, been criticized in that (lateral prefrontal) activation effects could stem from differential strategy use between groups, or less time spent on the task when performance is high^49^. Indeed, the memory training group (post-training) was asked to use the method of loci during memory encoding and order recognition, their strategy thus differed from participants in both control groups. Furthermore, we found that the memory training group (post-training) showed slower response times during order recognition, potentially related to increased memory search when mentally re-tracing previously studied information along the imagined path. Response time differences between the groups thus appear unlikely to have influenced activation decreases since the memory training group actually spent more time-on-task. Also, our results were directly related to memory performance at the four-months re-test, speaking for the relevant association of task-based activation decreases and behavioral improvements.

Moreover, durable memory formation relies on consolidation during rest that is thought to stabilize memory content. This entails communication between hippocampal-neocortical networks^25–27^, potentially reflecting “replay” of neuronal ensembles that were engaged during the preceding experience^29,30^. Across participants of the training study, we found increased hippocampal connectivity during rest after training with the lateral prefrontal cortex, left angular gyrus, parahippocampal regions, and the caudate nucleus that was higher the more durable memories were formed (**Fig. 5c**). Follow-up analyses revealed that these effects were specific to the memory training group after training but were not present in any of the control groups (**Fig. 5d**). Connectivity effects during consolidation were generally less widespread in the athlete compared to the training sample and were centered on increased hippocampal-cerebellar connectivity at higher memory performance. The cerebellum was associated with hippocampal-dependent navigation^50^ and might thus contribute to the consolidation of previously studied material. Due to their long-standing experience with the mnemonic technique, memory athletes (compared to participants of the training study) might have formed even stronger memories already during the tasks, thereby alleviating the need for additional consolidation during rest.

Altogether, we found that memory training enhanced durable memories. In both memory athletes and initially mnemonics-naïve participants after memory training, we found decreased brain activation in lateral prefrontal, as well as in posterior parahippocampal and retrosplenial cortices during encoding and recognition, respectively. These activation decreases were partly associated with better memory performance at a four-months follow-up, indicating that participants were able to successfully use the method even after several months. Importantly, these task-based effects were paralleled by increased hippocampal-neocortical connectivity during rest that was higher the more durable memories participants formed. Our findings are the first to demonstrate that mnemonic training boosts durable memory formation via decreased task-based activation and increased consolidation thereafter. In line with conceptual accounts of neural efficiency, this highlights a complex interplay between brain activation and connectivity critical for extraordinary memory.

## Methods

### Participants of athlete and training studies

Details about participant recruitment were described previously^3^. In brief, we tested 23 memory athletes (age: 28 ± 8.6 years, 9 female) that were ranked among the top 50 of the world’s memory sports (http://www.world-memory-statistics.com). These participants were compared to an equally sized control sample that was matched for age, sex, handedness, smoking status, and IQ, recruited among gifted students of academic foundations and members of the high-IQ society Mensa. Six participants of the matched control group were selected from the training study based on their cognitive abilities within the screening session (see below), evenly sampled from the three groups. Together, all participants were part of the so-called “athlete study”. Of the 23 memory athletes, 17 completed a word list encoding and order recognition task inside the MR scanner; current analyses were thus restricted to a sub-sample of participants (memory athletes: *N* = 17 (age: 25 ± 4 years, 8 female); matched controls: *N* = 16 (age: 25 ± 4 years, 7 female); see also **Methods, MRI data processing: task data** and **Methods, MRI data processing: resting-state periods** for a detailed description of exclusions).

Next, we recruited 51 male participants (age: 24 ± 3 years; all students at the University of Munich) to test the behavioral and neural effects of mnemonic training in a mnemonics-naïve participant sample (i.e., the so-called “training study”). Based on cognitive performance determined during an initial screening session (including fluid reasoning, memory abilities, see^3^), participants were pseudo-randomly assigned to three groups to ensure similar cognitive baseline levels between the groups. A first group of participants underwent a six-weeks training in the method of loci between the two test sessions (memory training group). These participants were directly compared to a sample who underwent a *n*-back working memory training between the sessions (active controls), and to a group who did not undergo any intervention (passive controls). Current analyses included 50 participants (memory training group, *N* = 17 (age: 24 ± 3 years); active controls, *N* = 16 (age: 24 ± 3 years); passive controls, *N* = 17 (age: 24 ± 4 years); see also **Methods, MRI data processing: task data** and **Methods, MRI data processing: resting-state periods** for a detailed description of exclusions). All participants provided written informed consent prior to participation and the study was reviewed and approved by the ethics committee of the Medical Faculty of the University of Munich (Munich, Germany).

### Study procedures and tasks

Participants of the memory athlete study completed a single MRI session (**Fig. 1a**). Participants of the training study took part in two MRI sessions that were placed six weeks apart, as well as in a behavioral session after four months (**Fig. 1b**). After the first MRI session (i.e., pre-training session), participants were pseudo-randomly grouped into one of three training groups and completed a training in the method of loci (memory training group), a working memory training (active controls), or no intervention (passive controls). Six weeks following the pre-training session, participants were invited to the second MRI session (i.e., post-training session), and were asked to complete a behavioral re-test four months thereafter.

#### Method of loci training

A detailed description of cognitive training procedures can be found here^3^. In brief, participants of the memory training group were familiarized with the method of loci at the Max Planck Institute of Psychiatry, after which they completed 30 min of training each day for 40 days at home. The training was completed and monitored using a web-based training platform (https://memocamp.com). Participants came into the laboratory for an interview (within small groups of 2-3 participants) regarding potential training problems once every week, they were trained under direct supervision, and received the training plan for the following week.

#### Working memory training

Participants of the active control group were familiarized with the dual *n*-back task where participants had to monitor and update a series of both visually presented locations and auditorily presented letters^51^. Participants completed 30 min of training each day for 40 days. The training was completed using a home-based working memory training program and training results were monitored daily. As above, the active controls came into the laboratory once a week for an interview (within small groups of 2-3 participants) regarding potential training problems and for a training under direct supervision. The passive controls did not receive any training between the two sessions.

#### General structure of MRI sessions

Each MRI session (athlete and training study) started out with the acquisition of a structural brain image, a baseline resting-state period, followed by the word list encoding and order recognition tasks, as well as a post-task resting-state period (**Fig. 1c**). Participants then performed a free recall test in the behavioral laboratory 20 min after exiting the MR scanner (i.e., immediate free recall), and another free recall test 24 hours later via phone interview (i.e., delayed free recall). Participants of the memory athlete study only performed the immediate but no delayed free recall test.

#### Resting-state periods

A first 8-minute resting-state period was acquired at the start of each MRI session (i.e., baseline rest; **Fig. 1c**). To assess intrinsic connectivity changes related to memory consolidation, another resting-state period (8 min) was placed after the order recognition task (i.e., post-task rest). Thus, participants of the memory athletes and training studies completed two and four resting-state periods, respectively.

#### Word list encoding task

A list of 72 concrete nouns was presented within each session. Words of both lists were counterbalanced for word length and frequency, were presented in random order, and the order of lists was balanced across participants. After an initial instruction (5 s), words were presented individually (2 s), separated by a jittered interval ranging between 2 and 5 s (mean = 3.5 s) during which a fixation cross was presented (**Fig. 2a**). Additionally, another fixation period (30 s) was inserted after every sixth word. Memory athletes and the memory training group (post-training) were asked to use the method of loci during word list encoding.

#### Order recognition task

During the order recognition task, participants viewed 24 triplets of words based on material from the previously encoded word list. A brief cue indicated the start of the next trial (2 s) after which a triplet was presented (10 s) and participates had to indicate if the word order was the same as presented before (3 s; answer options “same, sure”, “same, maybe”, “different, maybe”, and “different, sure”; **Fig. 3a**). Triplet presentations were separated by a control condition during which participants were asked if triplets that consisted of new words were shown in ascending or descending order according to their number of syllables. Memory athletes and the memory training group (post-training) were asked to use the method of loci during order recognition.

#### Free recall tests

Following MR scanning (approx. 20 min later), participants were asked to freely recall (i.e., to write down) the 72 words studied during the preceding word list encoding task (i.e., immediate free recall test). After 5 min, participants were asked if they would need more time, and the free recall test was terminated after an additional 5 min. Another free recall test (5+5 min) was performed via telephone 24 hours later (i.e., delayed free recall test). Performance was determined by the number of words correctly recalled, ignoring word order or spelling mistakes. Participants of the athlete study completed only the immediate but not the delayed free recall.

#### Re-test after four months

During the re-test four months after the post-training session, participants of the memory training study completed the word list encoding task once more, followed by a delay filled with a reasoning task (15 min), and a free recall task. All tasks were completed in the behavioral laboratory and the task material was the same as during the initial pre-training session. Participants of the memory athletes study and the memory training group (post-training) were asked to use the method of loci during word list encoding. Five participants of the memory training study (2 memory training group, 2 active controls, 1 passive controls) were not available for the re-test after four months.

#### Behavioral measures: memory durability

Memory durability was determined for participants of the memory training study by assessing performance at the immediate- (20 min) and delayed (24 hours) free recall test, for both the pre- and post-training session separately. This resulted in three types of responses (see also^21,52^): words that were (1) already forgotten during the immediate free recall test (“forgotten”), (2) recalled during the immediate but forgotten during the delayed free recall test (“weak”); or (3) recalled at both free recall tests (“durable”). Words that were not recalled at the immediate test but recalled at the delayed test were grouped together with words that were forgotten (number of words, mean ± SEM; (pre-/post-training) memory training group: 1.29 ± 0.65/0.59 ± 0.21, active controls: 0.94 ± 0.48/1.29 ± 0.57, passive controls: 1.35 ± 0.49/0.35 ± 0.19).

We aimed at identifying activation and connectivity profiles that were associated with durable memory formation and thus calculated a behavioral “memory durability score” for each participant (see also^52^). We divided the number of durable by the total number of recalled words (durable ∩ weak; i.e. the proportion of durable memories), thereby normalizing individual memory durability scores for general memory performance. We did this separately for the pre- and post-training session and included these values as a covariate in group-level analyses (see below). We did not determine memory durability for the athlete study as these participants only completed the immediate but not the delayed free recall test.

#### Behavioral measures: recognition performance (d-prime)

Recognition performance was quantified using *d*-prime scores. To accommodate hit rates of 1 and false alarm rates of 0 in memory athletes and the memory training group (post-training), we adjusted the individual hit and false alarm rates (*z*-scored) of all participants by adding 0.5 and then dividing by the number of signal and noise trials +1, respectively^53^. *D*-prime was calculated as the difference between these adjusted hit and false alarm rates [*z*(hits)-*z*(false alarms)], collapsing across the different confidence levels (“sure”, “maybe”) as memory athletes and participants of the memory training group (post-training) had very few “maybe” responses (number of “maybe” triplets, mean ± SEM; athlete study, memory athletes: 0.47 ± 0.1, matched controls: 4 ± 0.43; training study (pre-/post-training), memory training group: 6.1 ± 0.61/1.53 ± 0.54, active controls: 4.47 ± 0.85/2.36 ± 0.61, passive controls: 3.35 ± 0.74/2.65 ± 0.85). There were very few missed responses which were collapsed together with incorrect triplets (number of missed triplets, mean ± SEM; athlete study, memory athletes: 0.18 ± 0.1, matched controls: 0.29 ± 0.14; training study (pre-/post-training), memory training group: 0.59 ± 0.19/0.65 ± 0.17, active controls: 0.47 ± 0.17/0.29 ± 0.14, passive controls: 0.35 ± 0.12/0.35 ± 0.15).

#### Statistical analysis of behavioral measures

Analysis of all behavioral data was carried out using R (https://www.Rproject.org). The general free recall performance of participants in both studies was reported previously^3^. Here, we used a set of independent-samples *t*-tests and ANOVA models to analyze novel data regarding memory durability and order recognition performance (i.e., number of triplets correctly recognized, *d*-prime, and response times). Significant interaction effects were followed-up with post-hoc *t*-tests and were corrected for multiple comparisons (Bonferroni). Alpha was set to 0.05 throughout (two-tailed). Any exploratory analyses are explicitly described as such.

### Imaging parameters

All imaging data were collected at the Max Planck Institute of Psychiatry (Munich, Germany), using a 3T scanner (GE Discovery MR750, General Electric, USA) equipped with a 12-channel head coil. We acquired 192 T2*-weighted blood oxygenation level-dependent (BOLD) images during each resting-state period, using the following EPI sequence: repetition time (TR), 2.5 s; echo time (TE), 30 ms; 34 axial slices; interleaved acquisition; field of view (FOV), 240 × 240 mm; 64 × 64 matrix; slice thickness, 3 mm; 1 mm slice gap. During each task (i.e., word list encoding, order recognition), we obtained 292 T2*-weighted BOLD images with the following EPI sequence: TR, 2.5 s; TE, 30 ms; flip angle, 90°; 42 ascending axial slices; FOV, 240 × 240 mm; 64 × 64 matrix; slice thickness, 2 mm. The structural image was acquired with the following parameters: TR, 7.1 s; TE, 2.2 ms; flip angle, 12°; in-plane FOV, 240 mm; 320 × 320 × 128 matrix; slice thickness, 1.3 mm.

### MRI data processing: task data

#### MRI data preprocessing and participant exclusions

The fMRI data were processed using SPM8 (http://www.fil.ion.ucl.ac.uk/spm/) in combination with Matlab (The Mathworks, Natick, MA, USA). The first four volumes were excluded to allow for T1-equilibration. The remaining volumes were realigned to the mean image of each session (athlete study), or across sessions (training study). The structural scan was co-registered to the mean functional scan and segmented into grey matter, white matter, and cerebrospinal fluid using the “New Segmentation” algorithm. All images (functional and structural) were spatially normalized to the Montreal Neurological Institute (MNI) EPI template using Diffeomorphic Anatomical Registration Through Exponentiated Lie Algebra (DARTEL)^54^, and functional images were further smoothed with a 3D Gaussian kernel (8 mm full-width at half-maximum, FWHM).

Head motion (quantified as framewise displacement, FD)^55^ was similar across groups for word list encoding and order recognition tasks during the pre- and post-training sessions (**Supplementary Results 8**, **Supplementary Table S13**). We excluded one participant due to technical problems with the MR images (training study, active controls), and one participant due to strong motion (*FD* = 103.39; athlete study, matched controls; motion affected only the word list encoding task but the participant was excluded from all analyses). This left 50 participants within the training study (memory training group, *N* = 17; active controls, *N* = 16; passive controls, *N* = 17) and 33 participants within the athlete study (memory athletes, *N* = 17; matched controls, *N* = 16).

#### fMRI data modeling and statistical analysis: word list encoding task

Memory athletes as well as participants of the memory training group (post-training) showed free recall performance close to ceiling level. Thus, employing a common subsequent memory analysis would have led to an uneven distribution of trials across participants (i.e., most memory athletes might remember all words and forget none, whereas this might be different for participants of the control groups). Thus, we opted for an alternative approach and tested activation changes during word list encoding compared to an implicit baseline, with individual memory durability scores (see above) added as a covariate during statistical inference.

The BOLD response for all trials during the word list encoding task was modeled as a single task regressor, time-locked to the onset of each trial. Instructions were binned within a separate regressor of no interest. All events were estimated as a boxcar function with a duration of 3 s (encoding) or 5 s (instructions), and were convolved with the SPM default canonical hemodynamic response function (HRF). To account for noise due to head movement, we included the six realignment parameters, their first derivatives, and the squared first derivatives into the design matrix. A high-pass filter with a cutoff at 128 s was applied. For participants of the memory training study, both sessions (i.e., pre- and post-training) were combined into one first-level model. The task regressors were then contrasted against the implicit baseline.

We then tested activation changes between the groups and over time on a second level, applying pair-wise comparisons between the three groups. Specifically, we used three separate random-effects, mixed ANOVAs with group (i.e., memory training group vs. active controls; memory training group vs. Passive controls; and active vs. passive controls) as a between-, and session (pre-, post-training) as a within-subjects factor. Individual memory durability scores were added as a covariate (see above). Conditions were compared using post-hoc *t*-tests. Differential activation between memory athletes and matched controls was investigated with an independent-samples *t*-test and the number of words freely recalled was added as a covariate.

#### fMRI data modeling and statistical analysis: order recognition task

Following the rational above, we tested activation changes during order recognition compared to a control condition (i.e., syllable counting) rather than contrasting correct and incorrect trials. Recognition trials were modeled as a single task regressor, time-locked to the onset of each trial (i.e., the presentation of the triplet) and with the duration set until a button press occurred (thus, the duration was equal to the response time). Instructions were binned within a separate regressor of no interest (duration 2 s). The remaining modeling steps as well as the statistical inference were performed identical to the word list encoding task (see above). Individual recognition performance (i.e., *d*-prime scores) was added as a covariate during group-level analyses for both studies.

### MRI data processing: resting-state periods

#### MRI data preprocessing and participant exclusions

Data from resting-state periods was processed using the Functional Magnetic Resonance Imaging of the Brain (FMRIB) Software Library (FSL, v5.0.1, https://fsl.fmrib.ox.ac.uk/fsl/fslwiki/)^56^. As a first step, the structural scan was processed (using *fsl_anat*), re-oriented to the MNI standard space (*fslreorient2std*), bias-field corrected (FAST), and brain extracted (BET). The functional images were preprocessed using FEAT. We excluded the first four volumes to account for T1 equilibration, performed motion correction (MCFLIRT), spatial smoothing with a Gaussian kernel (5 mm FWHM), and aligned images to the bias-corrected, brain-extracted structural image (FLIRT) using boundary-based registration. The structural image was aligned with the MNI 152 EPI template using non-linear registration (FNIRT). After manual inspection of the registered images, we used independent component analysis (ICA) to automatically detect and remove subject-specific, motion-related artifacts (ICA-based strategy for Automatic Removal Of Motion Artifacts, ICA-AROMA, v0.3-beta)^57,58^.

Head motion (*FD*; see above) was similar across groups for all resting-state periods (baseline, post-task rest) during the pre- and post-training sessions (**Supplementary Results 8**, **Supplementary Table 13**. We excluded two participants due to technical problems with the MR images (both training study, active controls), and one participant due to strong motion during the resting-state period (see above; athlete study, matched controls). This left 49 participants within the training study (memory training group, *N* = 17; active controls, *N* = 15; passive controls, *N* = 17) and 33 participants within the athlete study (memory athletes, *N* = 17; matched controls, *N* = 16).

#### Seed-based hippocampal connectivity and statistical analysis

To test for hippocampal whole-brain connectivity related to consolidation, we placed a bilateral seed within the anatomical boundaries of the hippocampus (taken from the Automatic Anatomical Labeling atlas, AAL)^59^ and calculated connectivity by regressing the average hippocampal time course against all other voxel time courses in the brain, resulting in connectivity maps for each resting-state period (baseline and post-task rest for both pre-/post-training sessions). We then created difference maps (post-task minus baseline rest) that yielded connectivity increases within each session and submitted them to group-level analyses.

The athlete study comprised a single MRI session and connectivity increases (i.e., difference maps, post-task minus baseline) were analyzed using a linear regression with free recall performance (i.e., the number of words freely recalled 20 min post-MRI scanning) added as a covariate of interest. Data from the training study (i.e., two MRI sessions) was analyzed in three steps: we investigated hippocampal coupling (1) during the pre-training session (post-task minus baseline rest), (2) during the post-training session (post-task minus baseline rest), and (3) changes from pre- to post-training sessions ([post-task minus baseline rest]_post_ minus [post-task minus baseline rest]_pre_) using three separate linear regression models with memory durability added as a covariate of interest (i.e., (I-II) the session-specific proportion of durable memories formed, or (III) the change in memory durability from pre- to post-training sessions).

### Statistical thresholds for fMRI analyses and anatomical labeling

Throughout the manuscript, and unless stated otherwise, significance for all MRI analyses was assessed using cluster-inference with a cluster-defining threshold of *p* < 0.001 and a cluster-probability of *p* < 0.05 family-wise error (FWE) corrected for multiple comparisons. The corrected cluster size (i.e., the spatial extent of a cluster that is required in order to be labeled as significant) was calculated using the SPM extension “CorrClusTh.m” and the Newton-Raphson search method (script provided by Thomas Nichols, University of Warwick, United Kingdom, and Marko Wilke, University of Tübingen, Germany; http://www2.warwick.ac.uk/fac/sci/statistics/staff/academic-research/nichols/scripts/spm/).

Anatomical nomenclature for all tables was obtained from the Laboratory for Neuro Imaging (LONI) Brain Atlas (LBPA40, http://www.loni.usc.edu/atlases/)^60^.

## Data availability

All anonymized data and analysis code are available upon request in accordance with the requirements of the institute, the funding body, and the institutional ethics board.

## Acknowledgements

The authors would like to thank the International Association of Memory (IAM, https://www.iam-memory.org) for their support in recruiting memory athletes. This work was financially supported by the Netherlands Organization for Scientific Research (NWO), the Volkswagen Foundation, and the COST Action (CA18106).

## Author contributions

BNK, DR, KO, SK, MC, MD designed the study; BNK, PS, SW, MC, MD collected the data; ICW analyzed the data and wrote the first version of the manuscript; ICW, CL, MD edited the manuscript; AS, GF, CL, MC, MD supervised the study; ICW, BNK, PS, SW, DR, KO, SK, AS, GF, CL, MC, MD discussed and interpreted results and commented on the manuscript.

## Materials & Correspondence

Correspondence and material requests should be addressed to ICW and MD.

## Supplementary Information

### Supplementary Results

#### Supplementary Results 1: Memory durability across participants of the training study pre- and post-training

Memory durability during pre- and post-training sessions is depicted in **Supplementary Fig. S1**. Memory performance was analyzed using two mixed ANOVAs (separated for pre- and post-training sessions) including the within-subjects factor memory type (weak, durable, forgotten) and the between-subjects factor group (memory training group, active-, and passive controls). During the pre-training session, we found a significant main effect of memory type (*F*(2,141) = 25.79, *p* < 0.0001), indicating that the majority of words studied during the preceding word list encoding task were forgotten (weak > forgotten: *t*(141) = −6.51, *p* < 0.001; durable > forgotten: *t*(141) = −6.03, *p* < 0.001), while a similar amount of weak and durable memories were formed (*p* = 0.885). The three groups did not significantly differ in their memory performance (main effect of group, *p* > 0.999), and there was no memory type × group interaction (*p* = 0.601). Thus, there were no group differences in memory performance before training.

During the post-training session, results demonstrated a significant memory type × group interaction (*F*(4,141) = 20.24, *p* < 0.001; main effect of group: *p* > 0.999; main effect of memory type: *F*(2,141) = 3.8, *p* = 0.025). While post-hoc *t*-tests revealed a similar amount of weak memories for participants of all three groups, we found significantly more durable memories in the memory training group compared to both active (*t*(141) = 5.24, *p* < 0.0001) and passive controls (*t*(141) = 6.68, *p* < 0.0001). Furthermore, the memory training group forgot significantly less than the active (*t*(141) = −4.09, *p* = 0.002) and passive controls (*t*(141) = −5.06, *p* < 0.0001).

#### Supplementary Results 2: Region-of-interest (ROI) analysis of fMRI data during word list encoding across participants of the training study pre- and post-training

To further elucidate the decrease in encoding-related activation in the memory training group from pre- to post-training as compared to both control groups, we performed region-of-interest (ROI) analysis based on the significant peak cluster coordinates from the whole-brain analyses (session × group interaction effects; two separate full factorial designs: memory training group vs. active controls, memory training group vs. passive controls; see **Supplementary Tables S3** and **S4**). We placed a sphere (8 mm radius) around the peak coordinate of each significant cluster and extracted average parameter estimates. **Supplementary Fig. S2a** shows parameter estimates (arbitrary units, a.u.) for each peak coordinate during word list encoding for the memory training group and active controls during pre- and post-training sessions. **Supplementary Fig. S2b** displays data from the analysis comparing the memory training group with the passive controls. Overall, ROI-analyses once more highlighted decreased activation levels within the memory training group post-training as compared to pre-training, and compared to active as well as passive control groups.

#### Supplementary Results 3: Response times (RTs) during the order recognition task across participants of the athlete and training studies

Response times (RTs) during order recognition were first tested across participants of the athlete study (memory athletes, *N* = 17; matched controls, *N* = 16), using a mixed ANOVA with the within-subjects factor memory type (correct-“sure”, correct-“maybe”, incorrect) and the between-subjects factor group (memory athletes, matched controls). Results are shown in **Supplementary Fig. S3a** and revealed a significant main effect of memory type (*F*(2,61) = 17.9, *p* < 0.0001), indicating longer RTs during correct-“maybe” compared to correct-“sure” (*t*(61) = −5.11, *p* < 0.0001) or incorrect responses (*t*(61) = 5.52, *p* < 0.0001). There were no differences between the groups (*p* = 0.71) and no significant memory type × group interaction (*p* = 0.511).

Across participants of the training study, RTs were tested using two mixed ANOVA with the within-subjects factor memory type (correct-“sure”, correct-“maybe”, incorrect) and the between-subjects factor group (memory training group, active, and passive controls), separated for pre- and post-training sessions. Results are shown in **Supplementary Fig. S3c**. Before training, there was no significant difference between the groups (*p* = 0.663) and no memory type × group interaction (*p* = 0.495), but participants displayed longer RTs during correct-“maybe” compared to correct-“sure” (*t*(124) = −3.37, *p* = 0.003) or incorrect responses (*t*(124) = 4.85, *p* < 0.0001; main effect of memory type, F(2,124) = 11.56, *p* < 0.0001). After training, we found a significant main effect of group (*F*(2,98) = 5.96, *p* = 0.004), showing longer RTs within the memory training group compared to passive (*t*(98) = −3.47, *p* = 0.002) and active controls (trending significance; *p* = 0.056). There was no significant main effect of memory type (*p* = 0.202) and no significant memory type × group interaction (*p* = 0.208).

#### Supplementary Results 4: D-prime during order recognition across participants of the training study pre- and post-training

*D*-prime scores during pre- and post-training sessions are depicted in **Supplementary Fig. S3b**. *D*-prime was analyzed using a mixed ANOVA including the within-subjects factor session (pre-, post-training) and the between-subjects factor group (memory training group, active-, and passive controls). Results showed a significant main effect of session (*F*(1,94) = 5.82, *p* = 0.018), indicating generally higher *d*-prime scores during the post-compared to the pre-training session (*t*(94) = −2.56, *p* = 0.012). However, there was no significant main effect of group (*p* = 0.977) and no session × group interaction (*p* = 0.261).

#### Supplementary Results 5: ROI-analysis of fMRI data during order recognition across participants of the training study pre- and post-training

To further test the decrease in recognition-related activation in the memory training group from pre- to post-training as compared to the passive controls, we performed ROI analysis based on the significant peak cluster coordinates from the whole-brain analyses (session × group interaction effects; full factorial design: memory training group vs. passive controls; **Supplementary Table S7**; the comparison of memory training group and active controls yielded a significant main effect of session but no interaction; see **Supplementary Table S8**). We placed a sphere (8 mm radius) around the peak coordinate of each significant cluster and extracted average parameter estimates. **Supplementary Fig. S4** shows parameter estimates (arbitrary units, a.u.) for each peak coordinate during order recognition for the memory training group and passive controls during pre- and post-training sessions. Overall, ROI-analyses once more highlight decreased activation levels within the memory training group post-training as compared to pre-training, and compared to active as well as passive control groups.

#### Supplementary Results 6: Hippocampal connectivity in memory athletes compared to matched controls

Since memory athletes showed free recall performance (20 min post-MRI) close to ceiling level (**Fig. 1e**), we repeated the analysis of hippocampal connectivity changes during rest without the covariate of interest. Results revealed increased hippocampal connectivity with the right middle occipital gyrus across all participants of the training study from before to after the tasks (including both memory athletes and matched controls, *N* = 33; one-sample *t*-test, contrast difference map (post-task > baseline rest); *p* < 0.05, FWE-corrected at cluster level using a cluster-defining threshold of *p* < 0.001, cluster size = 68 voxels; MNI peak coordinate of global maximum, x = 24, y = −91, z = 7; *Z*-value = 4.1, 93 voxels).

Additionally, we tested whether our results regarding the consolidation-related increase in hippocampal-cerebellar coupling (as presented in the main manuscript) were driven by differences between memory athletes and matched controls. Indeed, we found increased hippocampal coupling with the cerebellum in memory athletes compared to matched controls during rest after the task (memory athletes, *N* = 17; matched controls, *N* = 16; independent-samples *t*-test, contrast difference map (post-task > baseline rest); *p* < 0.05, FWE-corrected at cluster level using a cluster-defining threshold of *p* < 0.001, cluster size = 56 voxels; MNI peak coordinate of global maximum, x = −10, y = −70, z = −49; *Z*-value = 3.98, 63 voxels). There were no significant results for the reverse contrast (matched controls > memory athletes). Thus, our results of increased hippocampal-cerebellar connectivity at higher free recall performance, which we presented in the main manuscript, were driven by connectivity changes in memory athletes and were not present in matched controls.

#### Supplementary Results 7: Hippocampal connectivity and association with memory durability in the three training groups (post-training)

To further investigate potential group differences in hippocampal resting-state connectivity associated with durable memory consolidation after training, we performed pair-wise comparisons (i.e., three separate independent-samples *t*-tests, contrast post-training difference map (post-task > baseline rest), memory durability (post-training) added as a covariate of interest).

First, comparing the memory training group to the active controls, we found that memory durability was associated with increased connectivity between the hippocampus and the left inferior temporal cortex (memory training group > active controls; *p* < 0.05, FWE-corrected at cluster level using a cluster-defining threshold of *p* < 0.001, cluster size = 32 voxels; MNI peak coordinate of global maximum, x = −56, y = −35, z = −24; *Z*-value = 4.03, 37 voxels; no significant results for the reverse contrast active controls > memory training group). Second, comparing the memory training group to the passive controls, we found that memory durability was associated with increased hippocampal coupling with the right inferior temporal cortex, bilateral insula, and dorsal anterior cingulate cortex (memory training group > passive controls, **Supplementary Table S11**; no significant results for the reverse contrast passive controls > memory training group). Third, and as expected, there were no significant differences between active and passive controls. Altogether, the above results indicated that the linear relationship between hippocampal connectivity and memory durability differed between the groups (i.e., memory training group vs. active/passive controls).

To follow-up on this, we tested the association between memory durability and hippocampal coupling separately for each group using one-sample *t*-tests, again with memory durability (post-training) added as a covariate of interest. Similar to the results presented in the main manuscript, we found increased hippocampal connectivity with the lateral prefrontal cortex, superior temporal gyrus, striatum, bilateral insula, and the dorsal anterior cingulate cortex that positively scaled with individual memory durability across participants of the memory training group (**Supplementary Table S11**). We did not find any significant hippocampal connectivity changes that negatively scaled with the covariate, and there was no (positive or negative) association between memory durability and hippocampal connectivity in the active or passive control groups.

To summarize, our results of increased hippocampal-neocortical connectivity at higher memory durability in the post-training session, which we presented in the main manuscript, appeared specific to connectivity changes in the memory training group and were not present in any of the control groups.

#### Supplementary Results 8: Head motion during task and resting-state periods

We assured that brain results were not affected by differences in scan-to-scan motion between sessions or groups and calculated the framewise displacement (FD)^1^ for every scan at time *t* by *FD*(*t*) = |Δ*d_x_*(*t*)| + |Δ*d_y_*(*t*)| + |Δ*d_z_*(*t*)| + *r*|*α*(*t*)| + *r*|*β*(*t*)| + *r*|*γ*(*t*)|, where (*dx*, *dy*, *dz*) is the translational-, and (*α*, *β*, *γ*) the rotational movement. One participant displayed abnormally strong head motion and was excluded from all further analyses (*FD* = 103.39; athlete study, matched controls). Remaining *FD* values are listed in **Supplementary Table S13**. There was no significant difference in *FD* between memory athletes and matched controls during any of the task- or resting-state periods (independent-sample *t*-tests; memory athletes, *N* = 17; matched controls, *N* = 16; word list encoding: *p* = 0.476; order recognition: *p* = 0.6; baseline rest: *p* = 0.405; post-task rest: *p* = 0.183).

Additionally, there was no difference in *FD* between the three groups of the training study (memory training group, *N* = 17; active controls, *N* = 16; passive controls, *N* = 17) during word list encoding (all following results stem from one-way ANOVAs, factor group (memory training group, active, passive controls), main effects of group are reported, pre-training: *p* = 0.722; post-training: *p* = 0.378), order recognition (pre-training: *p* = 0.893; post-training: *p* = 0.124), or during any of the resting-state periods (pre-training, baseline rest: *p* = 0.071; pre-training, post-task rest: *p* = 0.555; post-training, baseline rest: *p* = 0.607; post-training, post-task rest: *p* = 0.951). Thus, head motion was generally small and was comparable between the groups.

## Supplementary Figures

**Fig. S1.:**
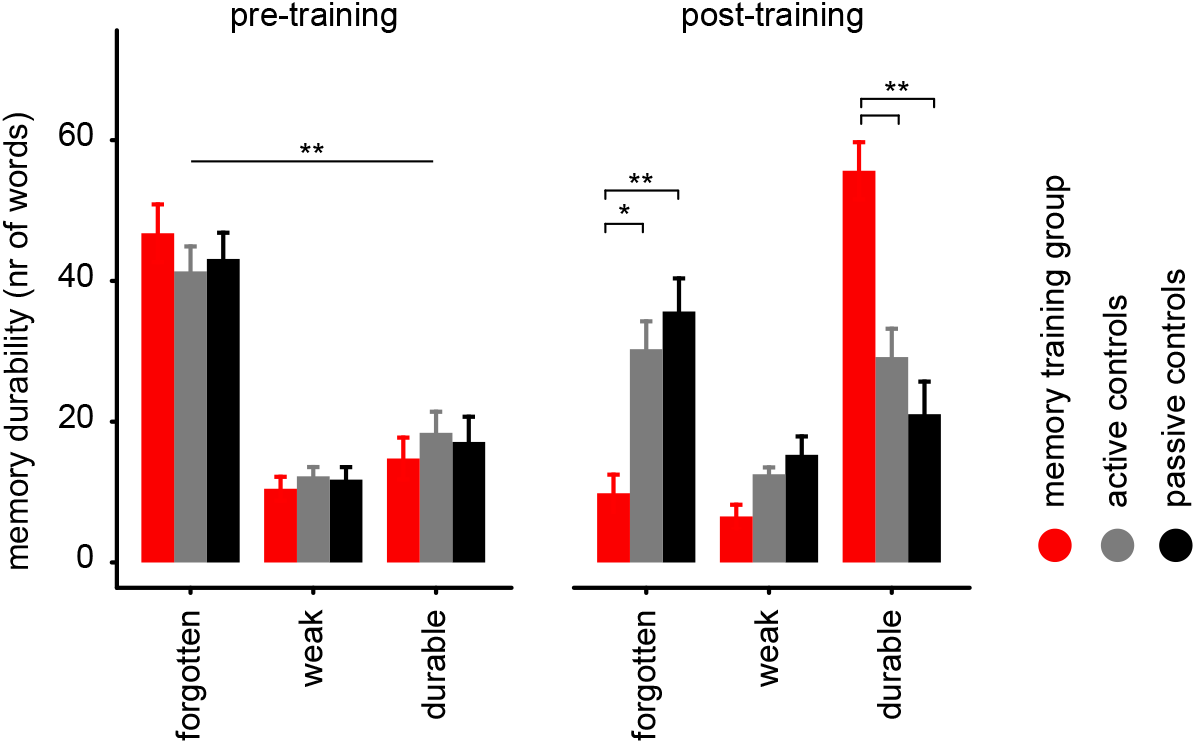
Memory durability across participants of the training study pre- and post-training. The number of words freely recalled that were forgotten/weak/durable) during pre- and post-training sessions. ** denotes *p* < 0.0001, * denotes *p* < 0.005. Error bars reflect the standard error of the mean, SEM.

**Fig. S2.:**
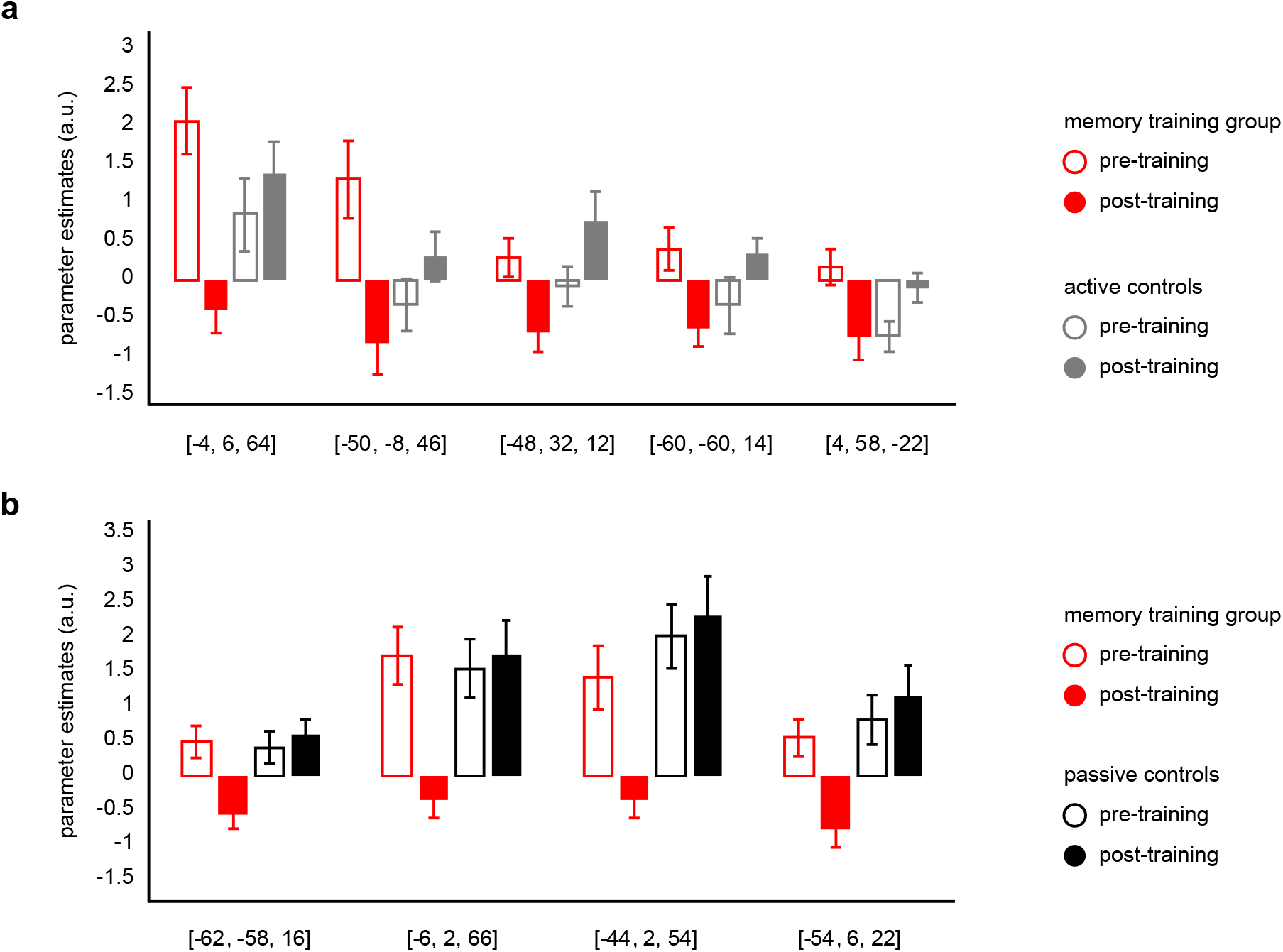
Region-of-interest (ROI) analysis of fMRI data during word list encoding across participants of the training study pre- and post-training. For visualization purposes, bar plots show extracted parameter estimates (arbitrary units, a.u.) from all significant clusters (8 mm sphere around MNI peak coordinates), stemming from whole-brain analyses during word list encoding (encoding > baseline). **(a)** Comparison of the memory training group with active controls (see also **Supplementary Table S3**). Clusters & anatomical labels: L superior frontal gyrus (x = −4, y = 6, z = 64), L precentral gyrus (x = −50, y = −8, z = 46), L inferior frontal gyrus (x = −48, y = 32, z = 12), L angular gyrus (x = −60, y = −60, z = 14), R superior frontal gyrus (x = 4, y = 58, z = −22). **(b)** Comparison of the memory training group with passive controls (see also **Supplementary Table S4**). Clusters & anatomical labels: L superior temporal gyrus (x = −62, y = −58, z = 16), L superior frontal gyrus (x = −6, y = 2, z = 66), L middle frontal gyrus (x = −44, y = 2, z = 54), L precentral gyrus (x = −54, y = 6, z = 22). L, left, R, right.

**Fig. S3.:**
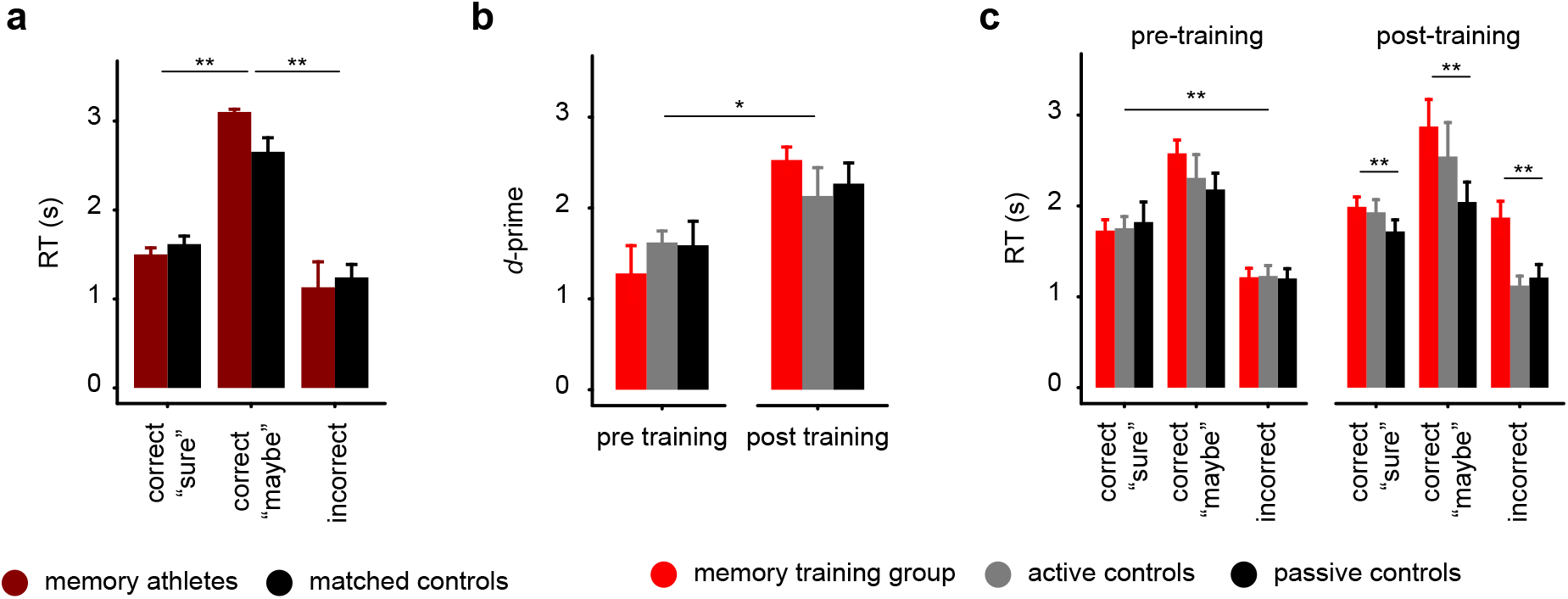
Response times (RT) and *d*-prime during the order recognition task across participants of the athlete and training studies. **(a)** Athlete study: response times (RT) in seconds (s) during correct-“sure”, correct-“maybe”, and incorrect responses for memory athletes and matched controls. ** denotes *p* < 0.0001. **(b+c)** Training study: **(b)** Recognition performance (*d*-prime) during pre- and post-training sessions. * denotes *p* < 0.05. **(c)** RT during correct-“sure”, correct-“maybe”, and incorrect responses during pre- and post-training sessions. ** denotes p < 0.0001, * denotes *p* < 0.005. Error bars reflect the standard error of the mean, SEM.

**Fig. S4.:**
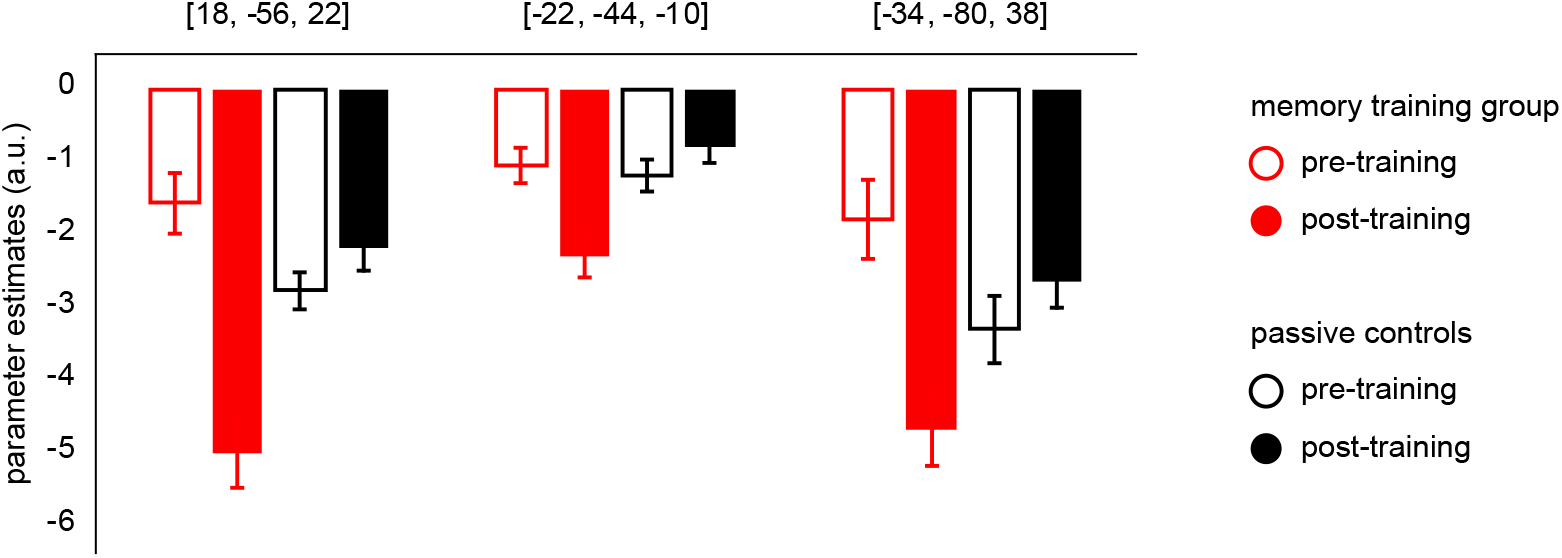
ROI-analysis of fMRI data during order recognition across participants of the training study pre- and post-training. For visualization purposes, bar plots show extracted parameter estimates (arbitrary units, a.u.) from all significant clusters (8 mm sphere around MNI peak coordinates), stemming from whole-brain analyses during order recognition (order recognition > baseline). Comparison of the memory training group with passive controls (see also **Supplementary Table S7**). Clusters & anatomical labels: R superior parietal gyrus (x = 18, y = −56, z = 22), L lingual gyrus (x = −22, y = −44, z = −10), L angular gyrus (x = −34, y = −80, z = 38). L, left, R, right.

## Supplementary Tables

### Supplementary Table S1

**Table S1:** Memory training enhances durable memory formation in initially mnemonics-naïve participants: results below present all significant pair-wise comparisons to follow-up on the significant group × memory type interaction (*F*(4,141) = 32.83, *p* < 0.0001; also indicated in **Fig. 1d**) and the main effect of memory type (*F*(2,141) = 13.27, *p* < 0.0001; main effect of group, *p* > 0.999).

| title of comparison | t-value | degrees of freedom | p-value |
| --- | --- | --- | --- |
| pair-wise comparisons between memory types: |  |  |  |
| weak > durable | -2.67 | 141 | 0.023 |
| weak > forgotten | 3.81 | 141 | < 0.001 |
| durable > forgotten | 6.48 | 141 | < 0.0001 |
| pair-wise comparisons, durable: |  |  |  |
| memory training group > active controls | 6.8 | 141 | < 0.0001 |
| memory training group > passive controls | 8.18 | 141 | < 0.0001 |
| pair-wise comparisons, forgotten: |  |  |  |
| memory training group > active controls | 5.88 | 141 | < 0.0001 |
| memory training group > passive controls | -6.6 | 141 | < 0.0001 |

### General note regarding tables displaying fMRI results

Significance for all MRI analyses was assessed using cluster-inference with a cluster-defining threshold of *p* < 0.001 and a cluster-probability of *p* < 0.05 family-wise error (FWE) corrected for multiple comparisons. The corrected cluster size (i.e., the spatial extent of a cluster that is required in order to be labeled as significant) was calculated using the SPM extension “CorrClusTh.m” and the Newton-Raphson search method (script provided by Thomas Nichols, University of Warwick, United Kingdom, and Marko Wilke, University of Tübingen, Germany; http://www2.warwick.ac.uk/fac/sci/statistics/staff/academic-research/nichols/scripts/spm/). MNI coordinated represent the location of peak voxels. We report the first local maximum within each cluster. Anatomical nomenclature for all tables was obtained from the Laboratory for Neuro Imaging (LONI) Brain Atlas (LBPA40, http://www.loni.usc.edu/atlases/; Shattuck et al., 2008). L, left; R, right; LH, left hemisphere; RH, right hemisphere.

### Supplementary Table S2

**Table S2:** Results from the word list encoding task, athlete study, differential activation between memory athletes and matched controls: independent-samples *t*-test, contrast encoding > baseline, covariate number of words freely recalled during the immediate test, critical cluster size: 125 voxels. Participant sample: memory athletes, *N* = 17; matched controls, *N* = 16. There were no significant effects for the reverse contrast.

| Contrast & brain region | MNI |  |  | Z value | Cluster size |
| --- | --- | --- | --- | --- | --- |
|  | x | y | z |  |  |
| *matched controls > memory athletes* |  |  |  |  |  |
| L inferior frontal gyrus | -48 | 32 | -8 | 4.55 | 226 |
| L inferior frontal gyrus | -46 | 20 | 16 | 4.38 | 370 |

### Supplementary Table S3

**Table S3:**
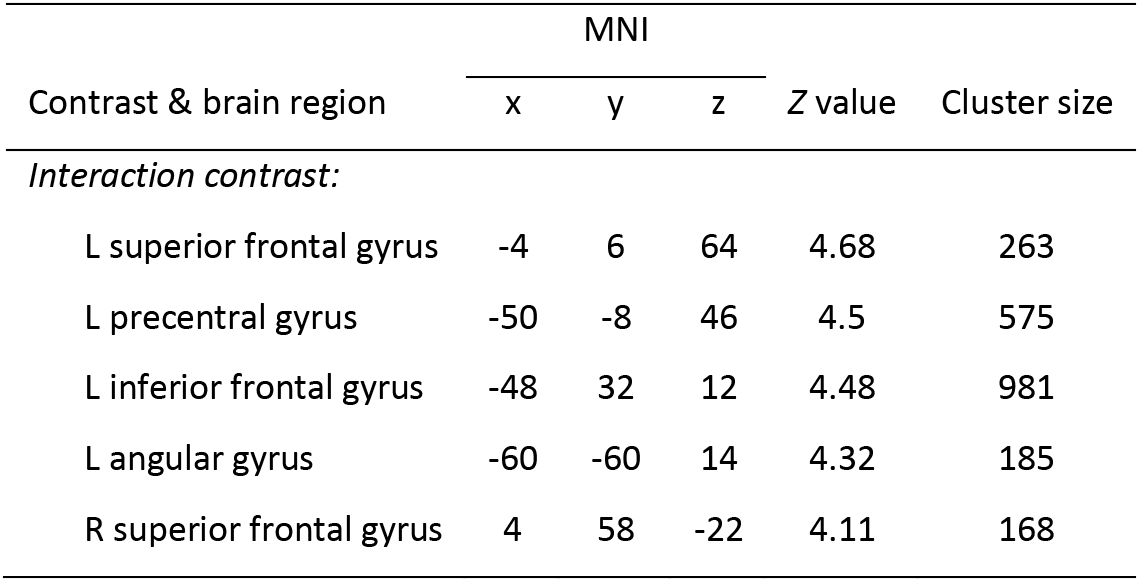
Results from the word list encoding task, training study, differential activation between the memory training group and active controls during pre- and post-training sessions: full factorial design with within-subjects factor session (pre-, post-training) and between-subjects factor group (active controls, memory training), contrast encoding > baseline, covariate memory durability score, critical cluster size: 138 voxels. Participant sample: memory training group, *N* = 17; active controls, *N* = 16. There were no significant results for the main effects of session or group. The interaction contrast shows the negative interaction of session × group; there were no significant effects for the reverse contrast.

### Supplementary Table S4

**Table S4:** Results from the word list encoding task, training study, differential activation between the memory training group and passive controls during pre- and post-training sessions: full factorial design with within-subjects factor session (pre-, post-training) and between-subjects factor group (passive controls, memory training), contrast encoding > baseline, covariate memory durability score, critical cluster size: 139 voxels. Participant sample: memory training group, *N* = 17; passive controls, *N* = 17. The interaction contrast shows the negative interaction of session × group. There were no significant results for the reverse contrasts.

| Contrast & brain region | MNI |  |  | Z value | Cluster size |
| --- | --- | --- | --- | --- | --- |
|  | x | y | z |  |  |
| *Main effect of session:* |  |  |  |  |  |
| *session 1 > session 2* |  |  |  |  |  |
| L inferior frontal gyrus | -46 | 22 | 14 | 4.64 | 687 |
| L superior frontal gyrus | -8 | 26 | 34 | 4.43 | 181 |
| L middle frontal gyrus | -36 | 0 | 54 | 4.41 | 396 |
| L angular gyrus | -48 | -64 | 22 | 4.02 | 422 |
| L superior frontal gyrus | -2 | 2 | 64 | 3.87 | 203 |
| R angular gyrus | 60 | -54 | 30 | 3.78 | 148 |
| *Main effect of group:* |  |  |  |  |  |
| *passive controls > memory training group* |  |  |  |  |  |
| L middle frontal gyrus | -52 | 8 | 46 | 4.76 | 1629 |
| R angular gyrus | 48 | -52 | 22 | 4.62 | 695 |
| R middle frontal gyrus | 42 | 58 | 18 | 4.6 | 1755 |
| L middle frontal gyrus | -32 | 52 | 24 | 4.57 | 502 |
| Cerebellum | 40 | -68 | -26 | 4.53 | 197 |
| Cerebellum | 46 | -62 | -30 | 4.4 | 254 |
| R superior frontal gyrus | 6 | 16 | 42 | 4.39 | 518 |
| R thalamus | 16 | -22 | 14 | 4.36 | 551 |
| L insular cortex | -28 | 16 | 8 | 4.31 | 221 |
| R precuneus | 4 | -56 | 50 | 4.3 | 783 |
| L angular gyrus | -38 | -60 | -42 | 4.18 | 427 |
| L angular gyrus | -48 | -56 | 18 | 4.14 | 335 |
| R insular cortex | 34 | 18 | 6 | 4.05 | 348 |
| R inferior frontal gyrus | 52 | 14 | 18 | 4.03 | 359 |
| *Interaction contrast:* |  |  |  |  |  |
| L superior temporal gyrus | -62 | -58 | 16 | 4.71 | 295 |
| L superior frontal gyrus | -6 | 2 | 66 | 4.52 | 323 |
| L middle frontal gyrus | -44 | 2 | 54 | 4.29 | 408 |
| L precentral gyrus | -54 | 6 | 22 | 4.11 | 531 |

### Supplementary Table S5

**Table S5:**
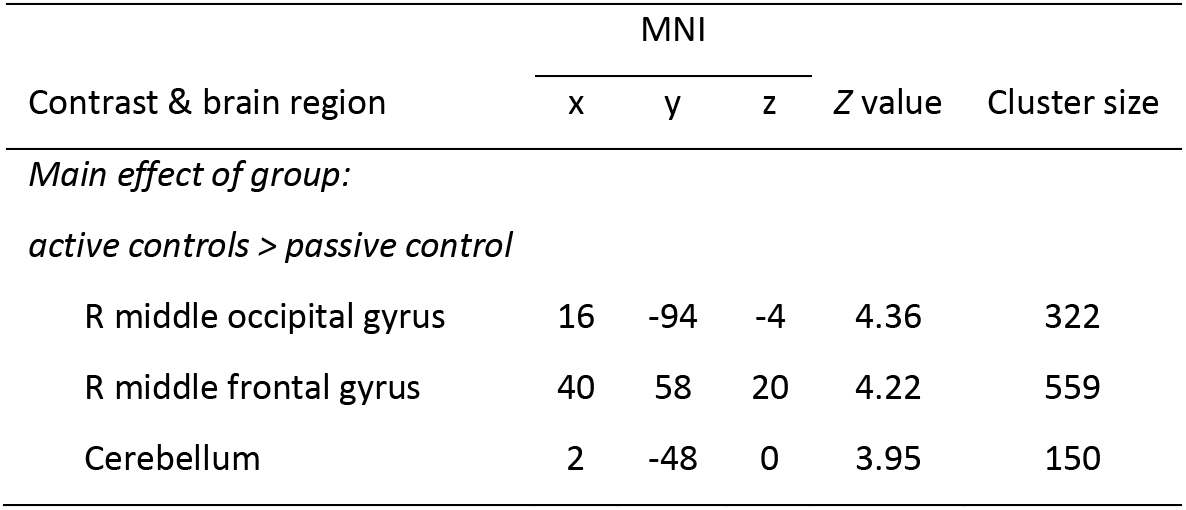
Results from the word list encoding task, training study, differential activation between active and passive controls during pre- and post-training sessions: full factorial design with within-subjects factor session (pre-, post-training) and between-subjects factor group (active, passive controls), contrast encoding > baseline, covariate memory durability score, critical cluster size: 140 voxels. Participant sample: active controls, *N* = 16; passive controls, *N* = 17. There were no significant results for the reverse contrast, no significant results for the main effect of session, or the session × group interaction.

### Supplementary Table S6

**Table S6:** Results from the order recognition task, athlete study, differential activation between memory athletes and matched controls: independent-samples *t*-test, contrast order recognition > baseline, covariate *d*-prime, critical cluster size: 123 voxels. Participant sample: memory athletes, *N* = 17; matched controls, *N* = 16. There were no significant effects for the reverse contrast.

| Contrast & brain region | MNI |  |  | Z value | Cluster size |
| --- | --- | --- | --- | --- | --- |
|  | x | y | z |  |  |
| *matched controls > memory athletes* |  |  |  |  |  |
| R superior parietal gyrus | 20 | -60 | 22 | 6.01 | 435 |
| L superior parietal gyrus | -16 | -58 | 23 | 5.31 | 501 |
| R parahippocampal gyrus | 30 | -40 | -12 | 4.54 | 142 |

### Supplementary Table S7

**Table S7:** Results from the order recognition task, training study, differential activation between the memory training group and passive controls during pre- and post-training sessions: full factorial design with within-subjects factor session (pre-, post-training) and between-subjects factor group (passive controls, memory training), contrast order recognition > baseline, covariate *d*-prime, critical cluster size: 127 voxels. Participant sample: memory training group, *N* = 17; passive controls, *N* = 17. There were no significant results for the main effects of session or group. The interaction contrast describes the negative session × group interaction; there were no significant results for the reverse contrast.

| Contrast & brain region | MNI |  |  | Z value | Cluster size |
| --- | --- | --- | --- | --- | --- |
|  | x | y | z |  |  |
| *Interaction:* |  |  |  |  |  |
| R superior parietal gyrus | 18 | -56 | 22 | 6.51 | 2568 |
| L lingual gyrus | -22 | -44 | -10 | 5.19 | 263 |
| L angular gyrus | -34 | -80 | 38 | 4.36 | 249 |

### Supplementary Table S8

**Table S8:** Results from the order recognition task, training study, differential activation between the memory training group and active controls during pre- and post-training sessions: full factorial design with within-subjects factor session (pre-, post-training) and between-subjects factor group (active controls, memory training), contrast order recognition > baseline, covariate *d*-prime, critical cluster size: 134 voxels. Participant sample: memory training group, *N* = 17; active controls, *N* = 16. There were no significant results for the reverse contrast. There were no significant result for the main effect of group or the session × group interaction.

| Contrast & brain region | MNI |  |  | Z value | Cluster size |
| --- | --- | --- | --- | --- | --- |
|  | x | y | z |  |  |
| *Main effect of session:* |  |  |  |  |  |
| *session 1 > session 2* |  |  |  |  |  |
| R superior parietal gyrus | 22 | -60 | 22 | 4.55 | 217 |
| L precuneus | -4 | -78 | 50 | 3.81 | 143 |
| L superior parietal gyrus | -12 | -56 | 16 | 3.72 | 148 |

### Supplementary Table S9

**Table S9:**
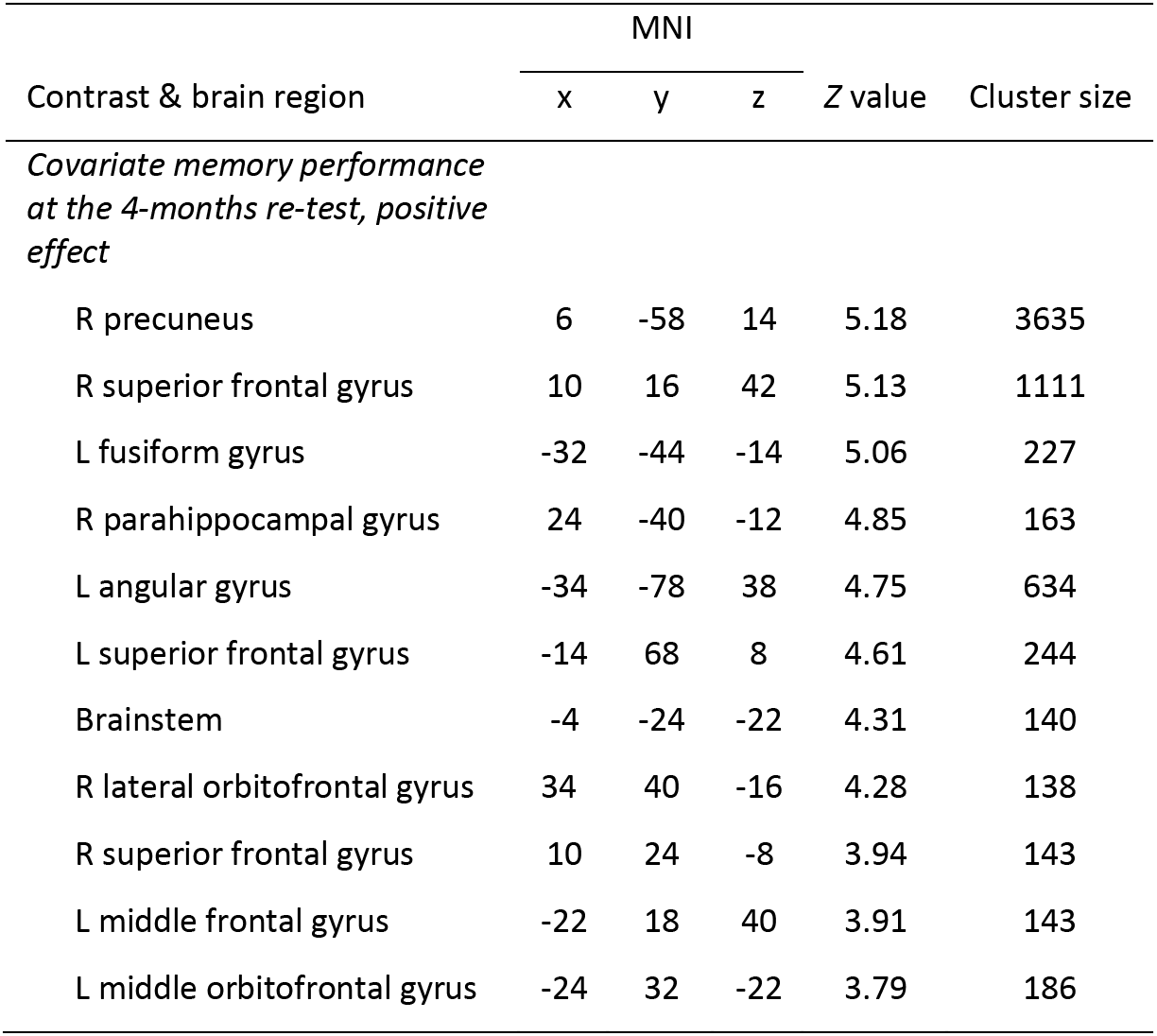
Activation decreases during order recognition and association with memory performance during the 4-months re-test: linear regression, contrast pre- > post-training session (order recognition > baseline), covariate change in memory performance (4-month re-test minus pre-training_20 min_), critical cluster size: 122 voxels. Participant sample: five subjects were not available for the re-test, analyses thus included 45 participants (memory training group, *N* = 16; active controls, *N* = 14; passive controls, *N* = 15). There was no negative association between activation decreases and the covariate.

### Supplementary Table S10

**Table S10:** Increased hippocampal-neocortical connectivity during post-training rest and relation with durable memory formation: linear regression, seed-based connectivity analysis (bilateral hippocampus seed), contrast post-task > baseline (post-training), covariate memory durability, critical cluster size: 35 voxels. Participant sample: memory training group, *N* = 17; active controls, *N* = 15; passive controls, *N* = 17. There were no effects for the reverse contrast, and no negative association between hippocampal connectivity and the covariate.

| Contrast & brain region | MNI |  |  | Z value | Cluster size |
| --- | --- | --- | --- | --- | --- |
|  | x | y | z |  |  |
| *Post-task > baseline rest* |  |  |  |  |  |
| R lingual gyrus | 7 | -63 | 7 | 5.98 | 2560 |
| L postcentral gyrus | -52 | -18 | 38 | 4.83 | 242 |
| R postcentral gyrus | 56 | -10 | 38 | 4.59 | 238 |
| L cingulate gyrus | 0 | 7 | 28 | 4.5 | 104 |
| L precentral gyrus | -4 | -28 | 70 | 4.19 | 185 |
| *Covariate memory durability, positive effect* |  |  |  |  |  |
| L parahippocampal gyrus | -32 | -32 | -14 | 5 | 84 |
| R inferior frontal gyrus | 42 | 18 | 0 | 4.44 | 70 |
| L inferior frontal gyrus | -49 | 35 | 10 | 4.44 | 108 |
| R caudate | 7 | 10 | -7 | 4.35 | 58 |
| L supramarginal gyrus | -52 | -38 | 24 | 4.12 | 40 |
| Cerebellum | 4 | -49 | -38 | 3.99 | 43 |
| Cingulate cortex | -14 | -18 | 28 | 3.97 | 41 |

### Supplementary Table S11

**Table S11:** Hippocampal connectivity and association with memory durability in the three training groups (post-training): For clarification, the analysis rationale is laid out once more. (1) Results from the comparison between the memory training group and active controls are described in the **Supplementary Results 7**. (2) Results from the comparison between the memory training group and passive controls are described in the table below (independent-samples *t*-test, contrast post-training difference map (post-task > baseline rest), memory durability (post-training) added as a covariate of interest, critical cluster size: 34 voxels. Participant sample: memory training group, *N* = 17; passive controls, *N* = 17. There were no significant results for the reverse contrast and no negative association with the covariate. (3) There were no significant differences between the active and passive control groups. (4) Significant results from (1) and (2) were followed-up with three separate one-sample *t*-tests, again with memory durability (post-training) added as a covariate of interest. We found hippocampal-neocortical coupling that positively scaled with memory durability in the memory training group (see table below, critical cluster size: 30 voxels, *N* = 17; see also **Fig. 5d**). There was no negative association with the covariate and no significant results in active or passive control groups.

| Contrast & brain region | MNI |  |  | Z value | Cluster size |
| --- | --- | --- | --- | --- | --- |
|  | x | y | z |  |  |
| *Memory training group > passive controls, covariate memory durability, positive effect* |  |  |  |  |  |
| L insular cortex | -38 | -18 | 4 | 4.39 | 105 |
| L superior frontal gyrus | -14 | -10 | 66 | 4.25 | 37 |
| L inferior occipital gyrus | -42 | -70 | -7 | 4.06 | 41 |
| R insular cortex | 35 | 10 | 7 | 3.78 | 37 |
| L superior frontal gyrus | -10 | -7 | 49 | 3.7 | 39 |
| *Memory training group, covariate memory durability, positive effect* |  |  |  |  |  |
| L superior frontal gyrus | -10 | -10 | 74 | 4.33 | 72 |
| L superior temporal gyrus | -60 | -35 | 10 | 4.15 | 65 |
| L superior frontal gyrus | -7 | 18 | 46 | 4.13 | 51 |
| L putamen | -28 | 4 | 7 | 3.95 | 98 |
| R inferior frontal gyrus | 38 | 18 | 4 | 3.93 | 36 |
| R precentral gyrus | 10 | -24 | 56 | 3.67 | 31 |

### Supplementary Table S12

**Table S12:** Hippocampal connectivity changes and association with memory durability from pre- to post-training: linear regression, seed-based connectivity analysis (bilateral hippocampus seed), contrast [post-task > baseline]_post_ > [post-task > baseline]_pre_, covariate memory durability, critical cluster size: 37 voxels. Participant sample: memory training group, *N* = 17; active controls, *N* = 15; passive controls, *N* = 17. There were no significant results for the reverse contrast, and no association with the covariate.

| Contrast & brain region | MNI |  |  | Z value | Cluster size |
| --- | --- | --- | --- | --- | --- |
|  | x | y | z |  |  |
| $[post\text{-}task > baseline\ rest]_{post}$<br>$> [post\text{-}task > baseline\ rest]_{pre}$ | | | | | |
| R lingual gyrus | 14 | -46 | 0 | 4.49 | 102 |
| R lingual gyrus | 4 | -80 | -7 | 4.02 | 49 |

### Supplementary Table S13

**Table S13:** Head motion during MR scanning: Head motion was quantified using framewise displacement (FD) separately for each task- and resting-state period, and for both pre- and post-training sessions. Values indicate average *FD* values per group (mean ± SEM). One participant (training study, active controls) displayed abnormally large head motion (*FD* = 103.39) during the word list encoding task and was excluded from all analyses (also not included in the table below). See sections **Methods, MRI data processing: task data** and **Methods, MRI data processing: resting-state periods** for further details regarding participant exclusions and final analysis samples.

| groups | single MRI session / pre-training MRI session |  |  |  | post-training MRI session |  |  |  |
| --- | --- | --- | --- | --- | --- | --- | --- | --- |
|  | baseline rest | word list encoding | order recognition | post-task rest | baseline rest | word list encoding | order recognition | post-task rest |
| *memory athletes* ($N = 17$) | 0.1 ($\pm 0.01$) | 0.1 ($\pm 0.01$) | 0.011 ($\pm 0.01$) | 0.09 ($\pm 0.01$) | | | | |
| *matched controls* ($N = 16$) | 0.11 ($\pm 0.01$) | 0.09 ($\pm 0.01$) | 0.11 ($\pm 0.01$) | 0.11 ($\pm 0.01$) | | | | |
| *memory training group* ($N = 17$) | 0.14 ($\pm 0.02$) | 0.11 ($\pm 0.01$) | 0.12 ($\pm 0.01$) | 0.12 ($\pm 0.01$) | 0.13 ($\pm 0.02$) | 0.15 ($\pm 0.02$) | 0.15 ($\pm 0.02$) | 0.13 ($\pm 0.02$) |
| *active controls* ($N = 16$) | 0.1 ($\pm 0.01$) | 0.1 ($\pm 0.01$) | 0.11 ($\pm 0.01$) | 0.11 ($\pm 0.01$) | 0.12 ($\pm 0.01$) | 0.12 ($\pm 0.01$) | 0.12 ($\pm 0.01$) | 0.13 ($\pm 0.02$) |
| *passive controls* ($N = 17$) | 0.12 ($\pm 0.01$) | 0.11 ($\pm 0.01$) | 0.11 ($\pm 0.01$) | 0.11 ($\pm 0.01$) | 0.13 ($\pm 0.01$) | 0.12 ($\pm 0.01$) | 0.13 ($\pm 0.01$) | 0.13 ($\pm 0.01$) |

